# Ciliary neurotrophic factor-mediated neuroprotection involves enhanced glycolysis and anabolism in degenerating mouse retinas

**DOI:** 10.1101/2021.12.21.473752

**Authors:** Kun Do Rhee, Yanjie Wang, Johanna ten Hoeve, Linsey Stiles, Thao Thi Thu Nguyen, Xiangmei Zhang, Laurent Vergnes, Karen Reue, Orian Shirihai, Dean Bok, Xian-Jie Yang

## Abstract

Ciliary neurotrophic factor (CNTF) acts as a potent neuroprotective cytokine in multiple models of retinal degeneration. To understand mechanisms underlying its broad neuroprotective effects, we have investigated the influence of CNTF on metabolism in a mouse model of photoreceptor degeneration. CNTF treatment improves the morphology of photoreceptor mitochondria, but also leads to reduced oxygen consumption and suppressed respiratory chain activities. Molecular analyses show elevated glycolytic pathway gene transcripts and active enzymes. Metabolomics analyses detect significantly higher levels of ATP and the energy currency phosphocreatine, elevated glycolytic pathway metabolites, increased TCA cycle metabolites, lipid biosynthetic pathway intermediates, nucleotides, and amino acids. Moreover, CNTF treatment restores the key antioxidant glutathione to the wild type level. Therefore, CNTF significantly impacts the metabolic status of degenerating retinas by promoting aerobic glycolysis and augmenting anabolic activities. These findings reveal cellular mechanisms underlying enhanced neuronal viability and suggest potential therapies for treating retinal degeneration.

## INTRODUCTION

Ciliary neurotrophic factor (CNTF) has long been recognized as a potent neuroprotective agent in the vertebrate retina [1]. Enhancement of neuronal survival by CNTF has been demonstrated in multiple animal models of retinal degeneration, ranging from zebrafish to canine [2]. Interestingly, CNTF is effective in rescuing photoreceptor degeneration due to various underlying causes, including mutations in photoreceptor-specific genes and damages induced by strong light or neurotoxins [3–11]. In addition to enhancing the viability of photoreceptors, CNTF has been shown to increase the survival of retinal ganglion cells and promote retinal ganglion cell axonal regeneration in optic nerve crush or transection models [12–19]. Based on its significant and broad neuroprotective effects for retinal neurons, an encapsulated cell implant producing a secreted form of recombinant human CNTF has been tested in clinical trials to treat hereditary and age-related retinal degenerative diseases [20–29]. The trials for retinitis pigmentosa (RP) with two CNTF doses for durations up to two years detected increased retinal thickness but did not onserve efficacy using the best corrected visual acuity as the primary outcome [25]. However, recent reports of the CNTF trial for treating macular telangienctasia (MacTel) type 2 have described morphological and visual function improvements using multiple readout parameters [30, 31]. The ongoing clinical trials also include treatment for glaucoma with retinal ganglion cell loss, a leading cause for blindness world-wide [32].

CNTF belongs to a subfamily of cytokines that share a tripartite receptor system including the two transmembrane receptors gp130 and LIFRβ and a ligand-specific alpha receptor (CNTFRα) [33, 34]. In developing and mature retinas, CNTF primarily stimulates the Jak-STAT and MEK-ERK signaling pathways [35–39]. Using cell type-specific gene ablation analysis in a mouse retinal degeneration model, we have shown that exogenous CNTF delivered to degenerating retinas initially activates STAT3 and ERK signaling in Muller glial cells through the gp130 receptor [40]. This initial event elicits an intercellular signaling cascade, which subsequently triggers gp130-mediated STAT3 activation in rod cells to promote photoreceptor survival [40]. Previous studies have also shown that the dosage and duration of CNTF treatment critically affect the rescued neurons, as a single dose injection of CNTF transiently affects the length of photoreceptor outer segments while prolonged exposure to high levels of CNTF can result in decreased visual function despite robust rescue of photoreceptors [7, 41–43]. Molecular analyses have revealed that CNTF-induced STAT3 phosphorylation followed by its dimerization and nuclear entry significantly influences the retinal transcriptome, resulting in rapid elevation of transcripts involved in innate immunity and growth factor signaling, as well as reduced expression of genes involved in phototransduction and maintenance of photoreceptor identities [44].

The mature retina is among the most metabolically active compartments in the central nervous system. Photoreceptors have high energy demands for regulating membrane potentials in response to visual cues [45, 46]. In addition, photoreceptors require continuous lipid and protein biosynthesis to sustain life-long outer segment renewal [47]. Both rod and cone photoreceptors primarily consume glucose, which is supplied by the choroidal vessels through the retinal pigment epithelium (RPE) [48–50]. Fatty acids can also be a fuel source available to photoreceptors through the blood supply and the catabolic process of outer segment degradation by the RPE [51–53]. Accumulating evidence indicate that cone photoreceptor survival relies on glucose availability, which is partially dependent on neighboring rod cells [54, 55]. Although mature photoreceptors contain a multitude of mitochondria distributed in the inner segments and at the synaptic termini, photoreceptors rely heavily on aerobic glycolysis under normal conditions [56–59]. Perturbing the glycolytic pathway regulatory enzymes phosphofructokinase (PFK) or lactate dehydrogenase (LDH) can result in deficits in outer segment renewal [58]. Ablation of genes encoding the glycolytic pathway enzymes hexokinase (HK2) in rod cells [60] or pyruvate kinase (PKM2) in rod or cone cells [61, 62] leads to photoreceptor dysfunction and degeneration in aging mice. Furthermore, protection of photoreceptor from strong light-induced damage involves AMPK activity, which is regulated by the cellular AMP to ATP ratio [63]. The cumulative data thus suggest that the cellular metabolic status of photoreceptors plays a crucial role in photoreceptor viability and function.

To elucidate cellular mechanisms underlying CNTF-dependent enhancement of neuronal viability, we have investigated the impact of CNTF signaling on retinal metabolism in a mouse model of retinitis pigmentosa, in which rod death precedes cone loss. By delivering the same secreted human CNTF used in clinical trials followed by molecular, cellular, and biochemical analyses, we demonstrate that CNTF treatment effectively influences the metabolic status of the degenerating retina, leading to elevated aerobic glycolysis and enhanced anabolism. These findings thus reveal a fundamental cellular mechanism by which a neurotrophic factor promotes neuronal viability in disease conditions.

## RESULTS

### CNTF treatment alters the morphology of rod photoreceptor mitochondria

To study the influence of CNTF on retinal metabolism under photoreceptor degeneration conditions, we used a mouse model of retinitis pigmentosa that expressed the dominant mutant pheripherin2 transgene Prph2(P216L) in the wild type photoreceptors [64]. This transgenic mouse model exhibits shortened photoreceptor inner segments, rudimental outer segments, and a relatively slow loss of photoreceptors, referred to as “retinal degeneration slow” (rds) herein. After mice reaching adulthood at postnatal day 25 (P25), the rds mutant retina contains approximately 80% of photoreceptors in number compared to the wild type. By P45, the loss of photoreceptors in the rds mutant has reached 50%. Subretinal delivery of a lentivirus expressing secreted form CNTF (LV-CNTF) at P25 halts the degeneration process, resulting in pan-retinal rescue of rod cells and partially restoration of the outer segment as well as the inner segment [40], where the majority of photoreceptor mitochondria reside.

To examine the influence of CNTF on rod mitochondria, we genetically labeled rod mitochondria by crossing Rho-iCre mice [65] with the PhAM mouse line that encodes a Cre-dependent mitochondrial targeting fluorescent reporter dendra2 [66]. The resulting mouse retina showed specific labeling of rod mitochondria with the dendra2 reporter in the inner segments and the ribbon synapses of wild type rod photoreceptors (Fig. 1a, 1b). The rds mutant retinas exhibited similar rod mitochondrial labeling, but with more ectopically located mitochondria within the outer nuclear layer where rod photoreceptor soma resided (Fig. 1a). To characterize the morphology of rod mitochondria, we performed super resolution structured illumination microscopy (SIM). The wild type rod mitochondria within the inner segment presented elongated cylindrical morphology, whereas those in the synaptic terminals were larger in size and granular in shape (Fig. 1c; Supplementary Video 1 and Video 2). In contrast, rod mitochondria in the rds mutant inner segment exhibited shortened and fragmental morphology (Fig. 1c; Supplementary Video 3). The LV-CNTF treatment of the rds retina resulted in elongation of the inner segments containing enlarged mitochondria both in the inner segments and in the synaptic termini (Fig. 1c; Supplementary Video 5), whereas control virus LV-IG treatment did not result in photoreceptor rescue or significant changes of mitochondrial morphology (Fig. 1c; Supplementary Video 4). TEM analysis also confirmed these morphological alterations detected by SIM (Fig. 1d).

**Figure 1.**
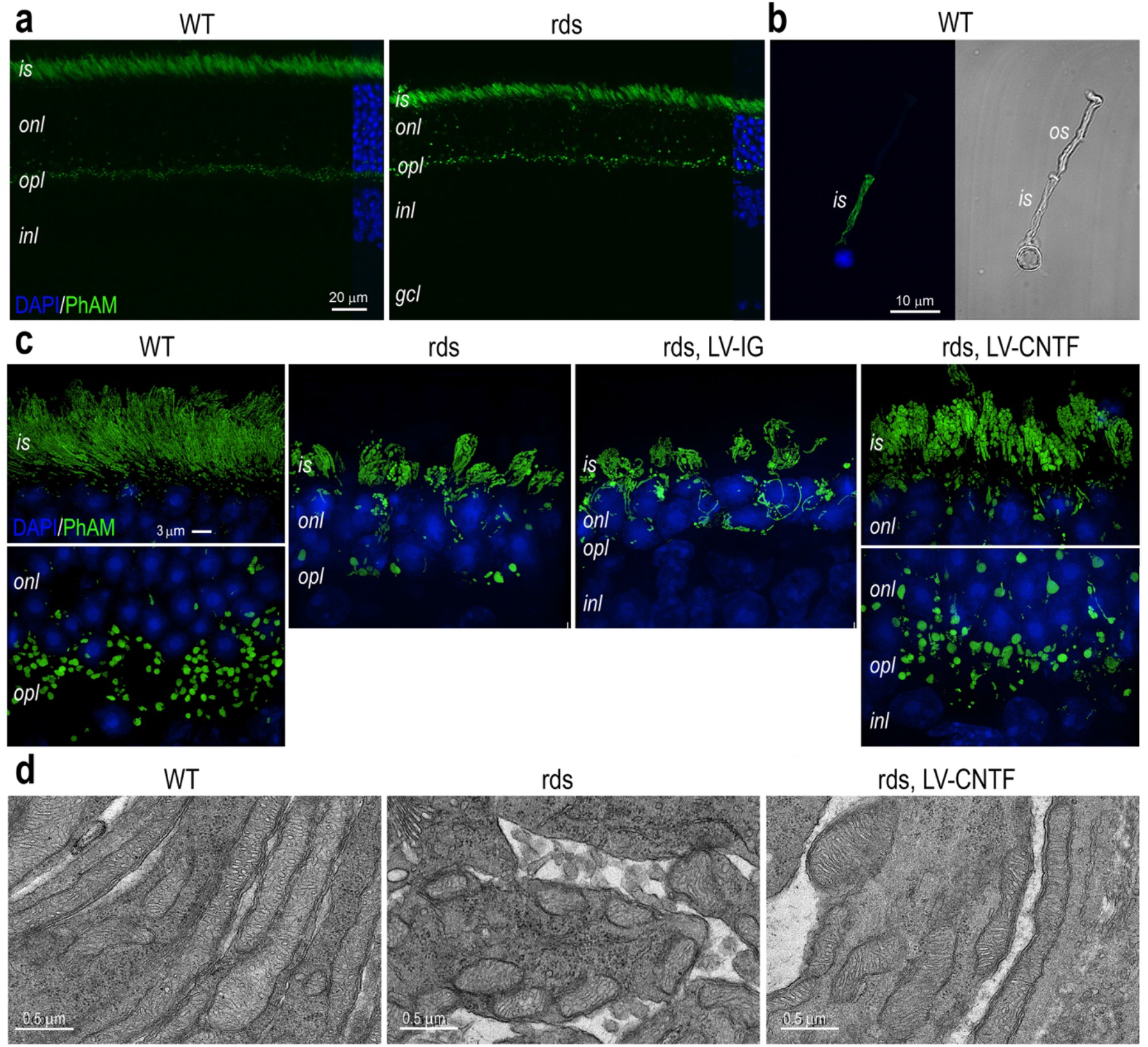
CNTF-induced mitochondrial morphological changes in rod photoreceptors. **a** Confocal microscopy images of P70 WT and rds retinas with the mitochondrial PhAM reporter activated in rod photoreceptors. **b** A dissociated P35 WT rod photoreceptor with PhAM reporter shown as Airyscan fluorescent image (left) and brightfield image (right). **c** SIM images show PhAM-labeled rod mitochondria in WT, rds, and rds mutant injected with LV-IG or LV-CNTF at P28 and harvested at P96. **d** TEM images of rod inner segments at P52 in WT, rds, and rds mutant injected with LV-CNTF. All image analyses used three independent retinas (N=3). Scale bars: a, 20 μm; b, 10 μm; c, 3 μm; d, 0.5 μm. WT, wild type; rds, Prph2 P216L mutant; is, inner segment; os, outer segment; onl, outer nuclear layer; opl, outer plexiform layer; inl, inner nuclear layer; gcl, ganglion cell layer.

Since mitochondria morphology may correlate with their functions [67], we characterized the morphological features of mitochondria in rod inner segments using acquired SIM imaging data (Fig.2; Source Data for Fig.2). The MitoMap analysis [68] confirmed that the rds mutant mitochondria showed marked departure from the wild type with increased sphericity and distribution isotrophy, but decreased compactness and surface to volume ratio (Fig. 2a). Principal component analysis indicated that in CNTF-treated rds retinas, morphological features of inner segment mitochondria displayed more resemblance to the wild type (Fig. 2b). However, in CNTF-treated rds retinas the mitochondrial population overall still exhibited significant differences from the wild type with regard to their compactness, surface to volume ratio, and distribution isotrophy (Fig. 2a).

**Figure 2.**
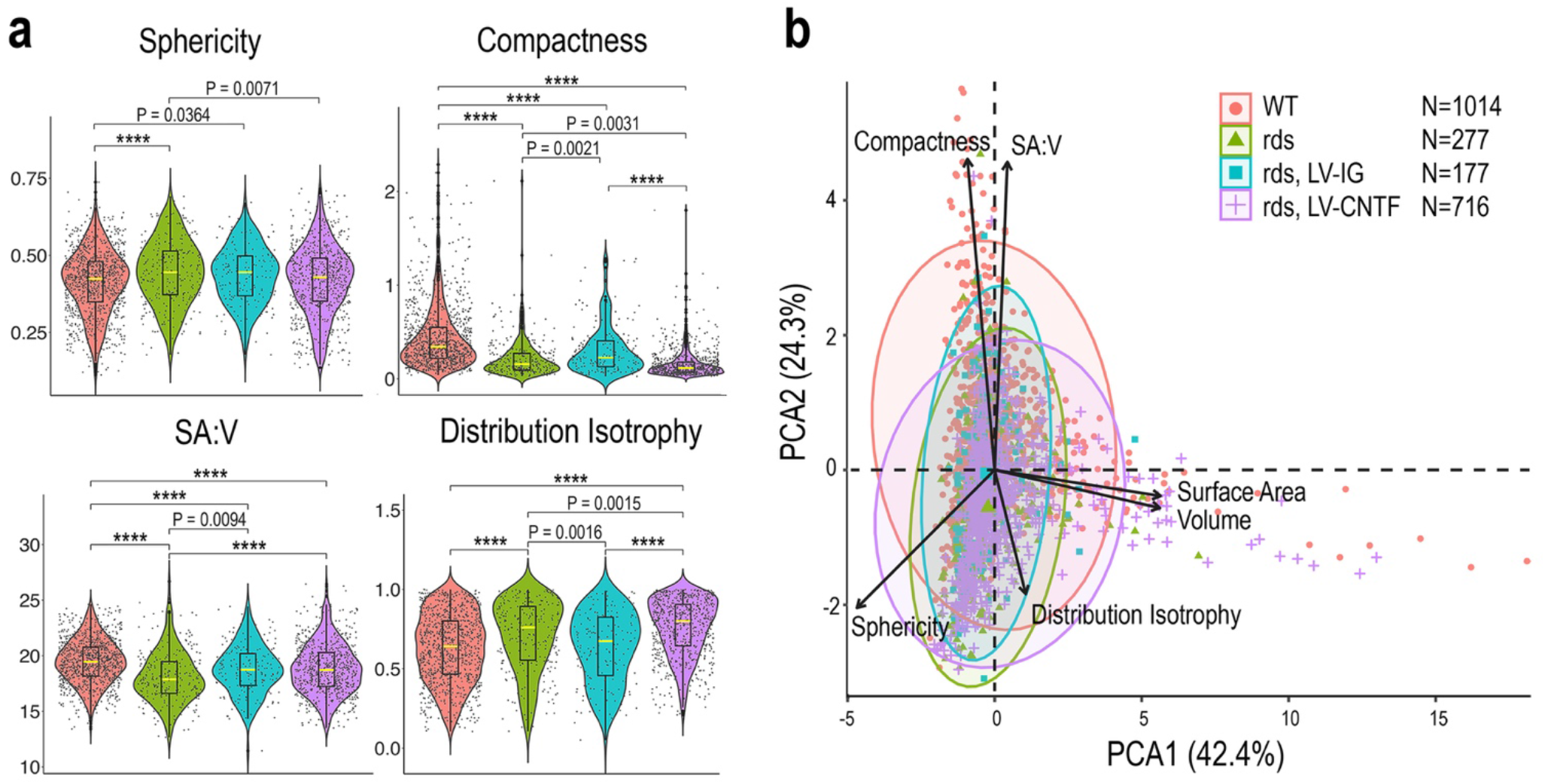
Characterization of rod photoreceptor mitochondrial morphological features. **a** Quantification of PhAM-labeled mitochondria in rod inner segments captured with SIM in WT, rds, and rds retinas (N=3) treated with LV-IG or LV-CNTF. Violin plots show geometrical features of individual mitochondrion, including sphericity, compactness, ratio of surface area to volume (SA:V), and distribution isotrophy. Dots represent individual mitochondria data points. Boxes represent 50% of the mitochondria, and yellow lines represent median values. Horizontal bounds of the box indicate 25th and 75th percentiles. The lower whiskers comprise the minimum data value within 1.5 times the interquartile range below the 25th percentile. The upper whiskers include the maximum value of the data within 1.5 times the interquartile range above the 75th percentile. One-way ANOVA and Tukey all-pairs test were applied with adjusted P values shown. P < 0.0001 is indicated as ****. **b** Principal component analysis (PCA) separates rod mitochondrial geometry. Separation in PCA1 is mainly driven by surface area, volume and sphericity, while compactness and SA:V contribute to separation in PCA2. N in b represent numbers of mitochondria used for the analysis shown in this figure.

### CNTF treatment affects mitochondrial respiration and suppresses respiratory complex activity

We next investigated the cellular respiration status using an Agilent XF96 Extracellular Flux Analyzer to simultaneously measure the oxygen consumption rate (OCR) and the extracellular acidification rate (ECAR), which may serve as a surrogate indicator for glycolysis (Fig. 3; Source Data for Fig. 3). Compared to the wild type, the rds mutant retina showed decreased basal OCR and significantly increased basal ECAR as early as P17 before overt degeneration of photoreceptors (Fig. 3a). Both the OCR reduction and ECAR elevation persisted through P35 during the continuous loss of rod cells in the outer nuclear layer. Despite the lengthening of rod inner segments in the rds retina, CNTF treatment did not result in an increased OCR, but significantly elevated ECAR (Fig. 3b), suggesting possible elevation of glycolysis. To examine whether CNTF affected oxidative phosphorylation, we first assayed the basal OCR, and then determined the amount of ATP-linked respiration and the maximal OCR by sequential additions of the ATP synthase inhibitor oligomycin followed by the proton gradient uncoupler FCCP (Fig. 3c). The OCR measurements indicated that CNTF treatment did not enhance the basal or maximal OCRs, but caused a reduction of ATP-linked OCR (Fig. 3c, 3d). These results suggested that CNTF treatment did not enhance mitochondrial respiratory chain-dependent oxygen consumption in the rds mutant retina.

**Figure 3.**
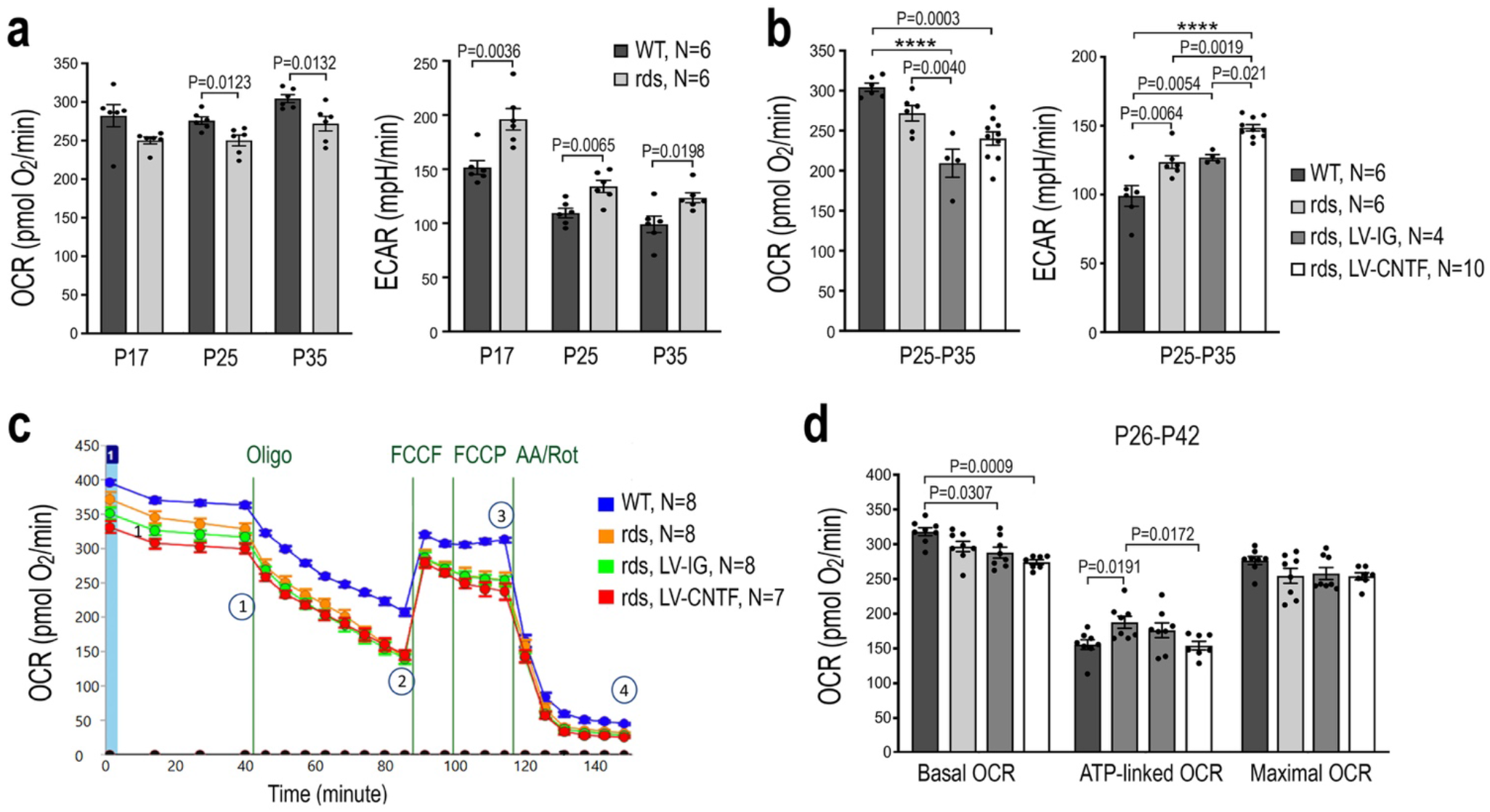
Influences on retinal cell respiration by the rds mutation and CNTF. **a** Seahorse metabolic flux analysis measuring basal OCR (left) and ECAR (right) for WT and rds retinas at P17, P25 and P35. For each age and genotype, N= 6 independent retinas were used. P values from two-tailed student *t* test are indicated. **b** Effects of CNTF on basal OCR and ECAR in rds retinas treated with LV-IG or LV-CNTF from P25-P35 compared to WT and non-treated rds retinas. Independent sample numbers N and adjusted P values from one-way ANOVA and Tukey all-pairs test are shown with P < 0.0001 indicated as ****. **c, d** Seahorse mitochondria stress tests to determine basal and maximum OCR, as well as ATP-linked OCR by sequential treatments with respiratory chain inhibitors oligomycin (oligo), uncoupling agent FCCP, antimycin A (AA) and rotenone (Rot) for WT, rds, and rds retinas treated with either LV-IG or LV-CNTF from P26-P42. **c** Seahorse assay tracings and the states indicating 1) basal OCR, 2) oligomycin inhibition of ATP synthase, 3) maximum OCR, and 4) total inhibition of respiration with AA and Rot. **d** Bar graphs show basal OCR, ATP-linked OCR, and maximal OCR. Independent sample numbers N and adjusted P values from two-way ANOVA and Tukey all-pairs test are shown. For **a**, **b**, **d**, data are presented as mean values ± SEM.

To validate the retinal tissue assay results, we analyzed mitochondrial respiration using isolated retinal mitochondria in the presence of different respiratory chain substrates and inhibitors (Fig. 4; Source Data for Fig. 4). In the presence of pyruvate and malate, substrates for complex I, mitochondria from rds retinas injected with the control virus LV-IG showed lower complex I-linked respiration compared to the wild type (Fig. 4a, 4b). LV-CNTF treatment of the rds retina caused a further reduction of complex I activity (Fig. 4a, 4b). When complex I was inhibited by rotenone and succinate was supplied, a slight reduction of complex II activity was detected in the rds retina (Fig. 4c, 4d). When complex I and complex III were both inhibited by rotenone and antimycin A to test complex IV activity, mitochondria from CNTF-treated rds retina showed a significant deficit when compared to the wild type (Fig. 4e, 4f). Together, these results demonstrated that the rds retina had an impaired mitochondrial respiratory chain, and CNTF treatment did not improve but instead caused a further suppression of mitochondrial respiratory chain function in the rds mutant.

**Figure 4.**
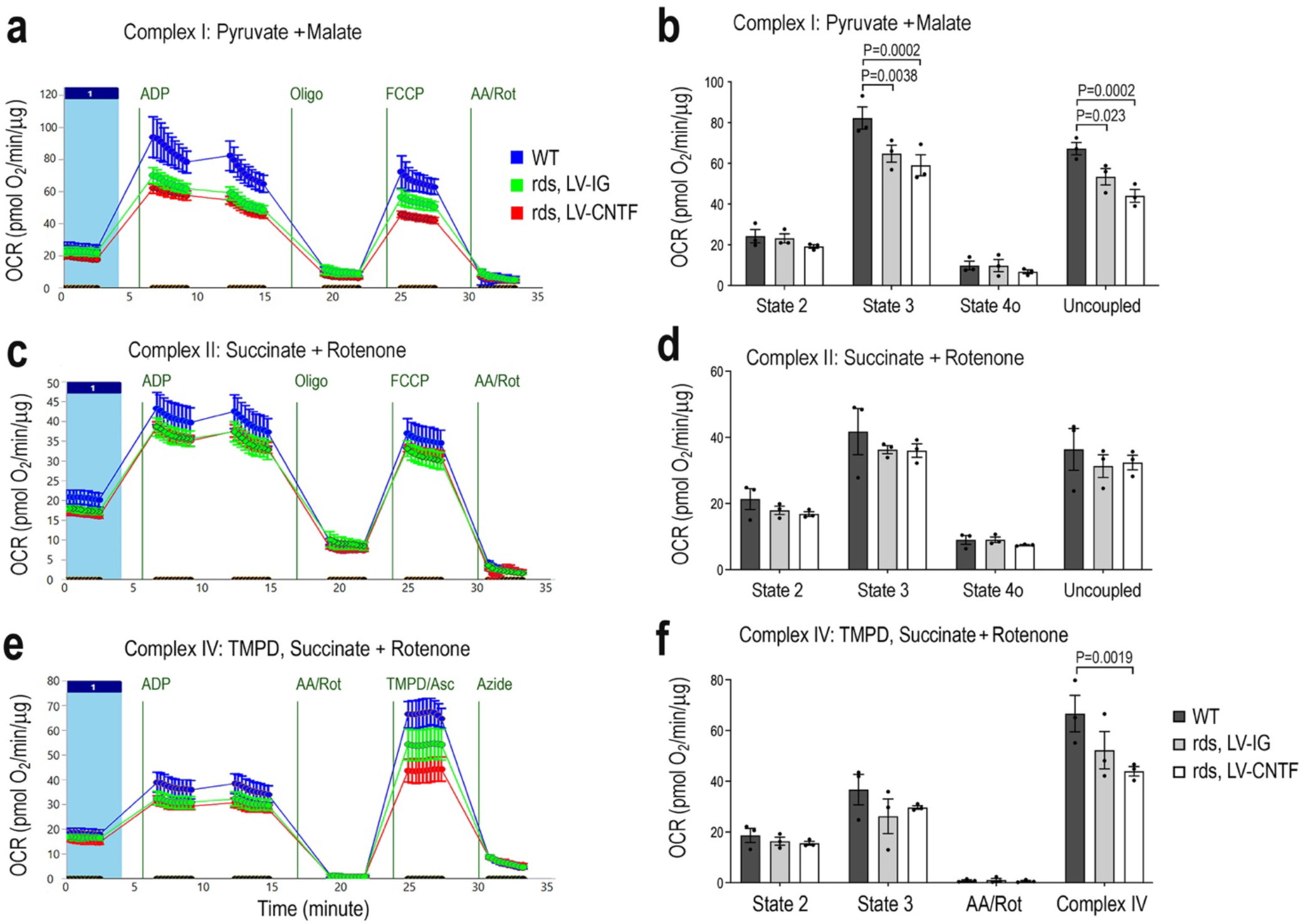
Quantification of mitochondrial respiratory chain complex activities. Mitochondria stress test using isolated mitochondria to examine activity of each complex from WT and rds retinas treated with LV-IG or LV-CNTF from P25-P49. OCR (pmole/min/μg protein) were measured. **a, b** Complex I respirometry tracing and bar graph using pyruvate and malate as substrates. **c, d** Complex II-driven respiration measured in the presence of succinate and complex I inhibitor rotenone. **e, f** Complex IV state 3 respiration was measured in the presence of succinate and rotenone, following inhibition of complex I and III, and injection of TMPD/ascorbate to measure Complex IV activity. Independent samples N=3, and each contains mitochondria from 4 retinas for all conditions. For **b**, **d**, **f**, data are presented as mean ± SEM. Adjusted P values derived from two-way ANOVA and Tukey all-pairs test are indicated.

### CNTF elevates energy production, aerobic glycolysis, and anabolic metabolites

To obtain a comprehensive assessment of the retinal metabolic status, we performed metabolomics analysis for wild type, rds, and rds retinas treated with either the control LV-IG or LV-CNTF virus. Quantification of cellular metabolites with glucose as a fuel revealed a dramatic impact of CNTF signaling on retinal metabolism (Supplementary Fig.1; Source Data of Metabolomics). Compared to the wild type, the rds mutant retina had a lower level of ATP whereas CNTF-treated rds retinas contained 1.5-fold of wild type level of ATP, and consequently a lower ADP/ATP ratio (Fig. 5a; Supplementary Fig. 2; Source Data for Fig. 5). In addition, the energy currency phosphocreatine (p-creatine), which is normally present at high concentrations in the brain and muscles to act as an energy buffer to quickly regenerate ATP [69, 70], was increased to 3.9-fold of the wild type level in CNTF-treated rds retinas (Fig. 5a). Furthermore, CNTF restored the level of GTP in the rds retina, which was reduced to 45% of the wild type level, and elevated 5’-methylthioadenosine to 1.4-fold of the wild type level (Fig. 5b; Supplementary Fig. 4). Consistent with the notion that CNTF signaling promoted aerobic glycolysis, we detected increases of glycolytic pathway intermediates fructose1,6 bisphosphate (F1,6BP), 3-phosphoglycerate (3PG) and phosphoenolpyruvic acid (PEP) (Fig. 5c; Supplementary Fig. 2). Furthermore, metabolomics analysis detected significant elevation of TCA cycle products, including aconitate, citrate, alpha ketoglutarate, and malate (Fig. 5d; Supplementary Fig. 2). Importantly, CNTF treatment also altered the reduction-oxidation status by fully restoring the antioxidant glutathione (GSH) in the rds retina from 50% reduction to the wild type level (Fig. 5e; Supplementary Fig. 2).

**Figure 5.**
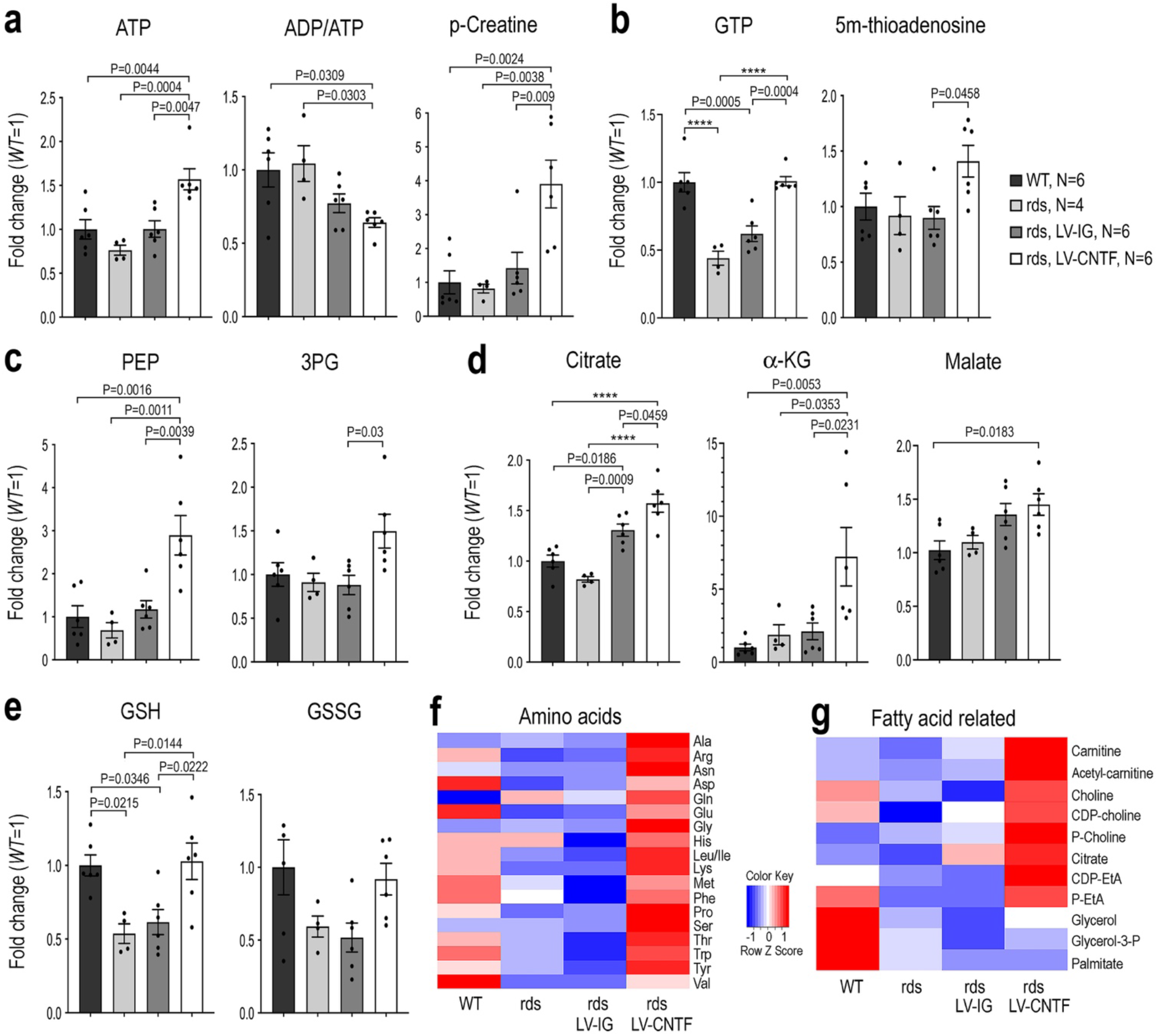
CNTF-dependent enhancement of aerobic glycolysis and anabolism. Metabolomics analysis of WT, rds, and rds mutant retinas treated with LV-IG or LV-CNTF from P25-P36 using glucose as a nutrient. **a** Energy metabolites ATP, phosphocreatine (p-creatine), and ADP/ATP ratios. **b** Nucleotide derivatives. **c** Glycolytic pathway intermediates 3-phosphoglycerate (3PG) and 2-phosphoenol pyruvate (PEP). **d** TCA cycle metabolites citrate, alpha-ketoglutarate (α-KG), and malate. **e** Redox related metabolites glutathione (GSH) and glutathione bisulfide (GSSG). **f** Heatmap shows retinal amino acid contents. **g** Heatmap shows fatty acid metabolic intermediates. For **a**-**e**, data are presented as mean ± SEM. Independent retinal sample numbers (N) and adjusted P values based on one-way ANOVA and Tukey all-pairs test are shown with P < 0.0001 indicated as ****.

Quantitative metabolomics analysis revealed that rds retinas treated with LV-CNTF, but not the control LV-IG virus, elevated cellular contents for the majority of amino acids to above the wild type levels (Fig. 5f; Supplementary Fig. 3). Similarly, multiple fatty acid biosynthetic intermediates that showed reduced levels in the rds retina were also elevated by CNTF treatment (Fig. 5g; Supplementary Fig. 2). The results of metabolomics analysis therefore indicated that CNTF signaling asserted a strong influence on retinal metabolism by promoting glycolysis, increasing energy supply, and enhancing anabolism in the rds mutant retina.

### CNTF signaling impacts expression of metabolic genes and enzymatic activities

To evaluate whether CNTF signaling influenced metabolism at the transcription level, we performed high throughput RNA-sequencing and analyzed retinal transcriptome of the rds mutant retinas treated with either the control LV-IG or LV-CNTF from P25 to P35 [44]. Compared to the wild type, LV-CNTF treatment led to increased expression of most glycolytic pathway transcripts, whereas LV-IG injection induced moderate gene expression elevation (Fig. 6a; Source Data for Fig. 6). CNTF treatment also elevated expression of many genes encoding the TCA cycle enzymes (Fig. 6b). Some of the genes involved in mitochondria stress and turnover, such as Pink1 [71], was elevated in the rds mutant (Fig. 6c). CNTF treatment suppressed the expression of Pink1 in the rds mutant, and elevated expression of key mitochondrial transcription factor Tfam [72], suggesting that CNTF signaling influenced mitochondrial dynamics. Transcriptome analysis also revealed that most nuclear encoded respiratory complex genes were up-regulated after CNTF treatment; however, several components for complex I, complex IV, and ATP synthase were downregulated compared to LV-IG treated rds retinas (Fig. 6d).

**Figure 6.**
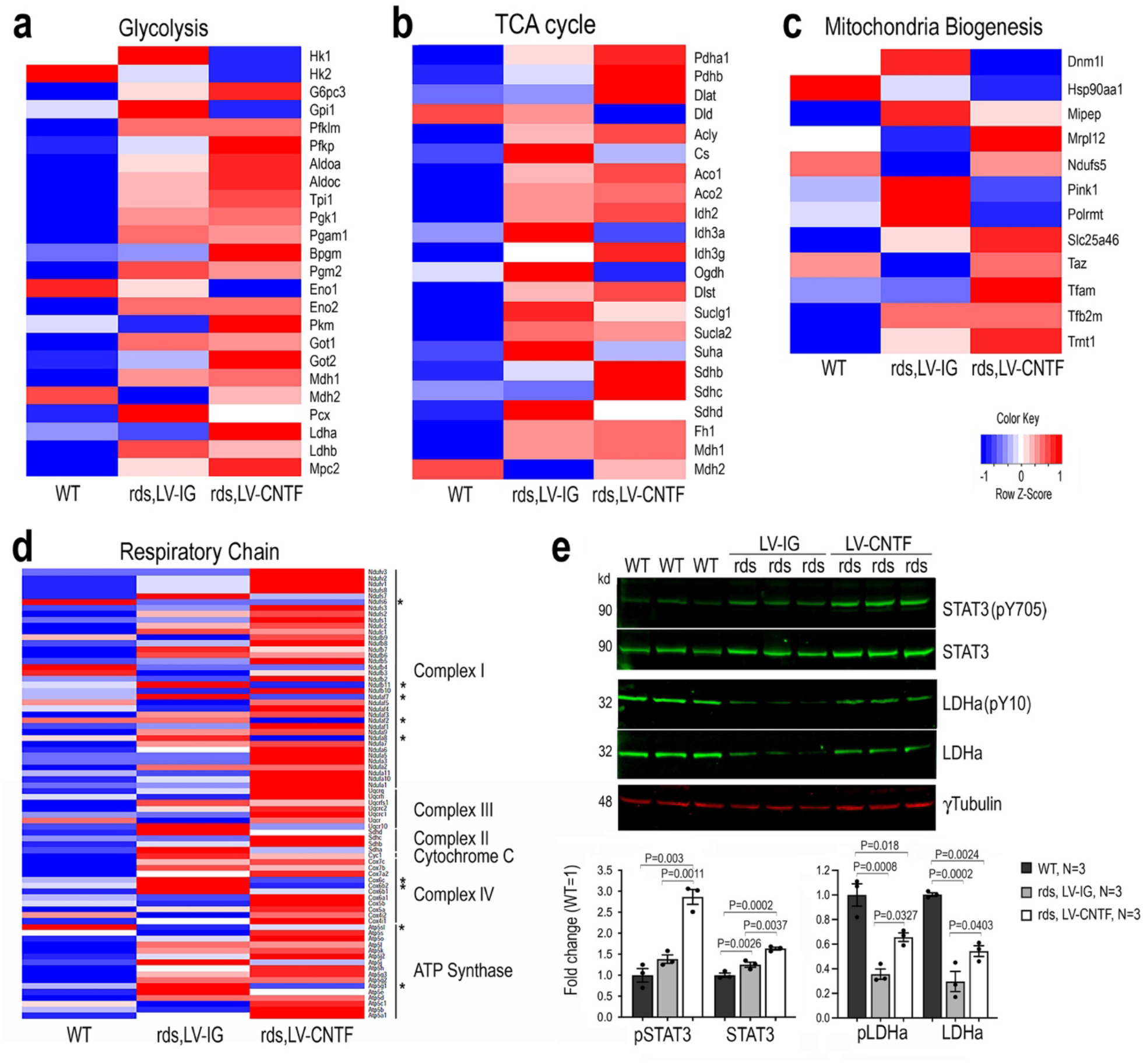
Influence of CNTF on metabolic gene expression and enzyme activity. **a-d** Transcriptome analysis of WT retina and rds retinas treated with LV-IG or LV-CNTF from P25-P35. Heatmaps show relative transcript levels (average of N=3, each N contains 2 retinas). **a**. Glycolytic pathway enzymes and Mpc2. **b** Mitochondrial Pdha1, Pdhb, and TCA cycle enzymes. **c** Participants of mitochondrial biogenesis. **d** Nuclear genome encoded mitochondrial respiratory chain components. Asterisks indicate genes with decreased transcripts compared to WT. **e** Western blot analysis of retinal extracts from WT retina and rds retinas treated with LV-IG or LV-CNTF from P25-P100. Bar graphs show quantification of CNTF signaling effector pY705 STAT3 and total STAT3, the active pY10 LDHa and LDHa. For **e**, data are presented as mean ± SEM. Independent retinal samples (N=3) and adjusted P values based on one-way ANOVA and Tukey all-pairs test are indicated.

Next, we performed Western blot analysis to examine the Jak-STAT signaling and the status of enzymes in the glycolytic pathway. As expected, CNTF treatment resulted in increased phosphorylation of STAT3 at Y705 as well as elevated total STAT3 protein (Fig. 6e; Source Data for Fig. 6). Consistent with the observed CNTF-induced elevation of LDHa mRNA (Fig. 6a), the active form of LDHa with Y10 phosphorylation were increased to near 2-fold of LV-IG treated rds retinas. These data thus further validated that CNTF signaling enhanced glycolysis in rds retinas.

### Contribution of glycolysis and OXPHOS in wild type and degenerating retinas

Since the metabolic changes in degenerative retinas were not well characterized, we examined the metabolic contribution from aerobic glycolysis and mitochondrial respiration in the wild type and rds retinas. Metabolomics analyses were performed under conditions with various metabolic inhibitors (Fig. 7a; Source Data for Fig. 7). In the presence of the complex IV inhibitor sodium azide (Supplementary Fig. 5), amino acid contents in both the wild type and CNTF-treated rds mutant retinas were significantly reduced (Fig. 7b; Supplementary Fig.6), indicating that the respiratory chain activities impacted cellular amino acid pools. Inhibition of complex IV also decreased CNTF-dependent elevation of lipid biosynthetic pathway intermediates (Fig. 7c). Quantification of ATP levels revealed that in the wild type retina, 87% of ATP production relied on oxidative phosphorylation (OXPHOS), whereas in the rds mutant only 65% of ATP production was mitochondrial dependent (Fig. 7e), suggesting that the degeneration condition itself had led to decreased dependency on OXPHOS. In CNTF-treated rds mutant, azide caused 62% reduction of ATP, indicating that a significant portion of the total ATP, about 50% of the wild type level, was not generated through mitochondrial respiration (Fig. 7e). Inhibition of complex IV by azide resulted in 80% reduction of acetyl-CoA in the wild type, but 66% reduction of acetyl-CoA and malate in CNTF treated rds retinas (Fig. 7f, 7g). Moreover, analysis using azide showed that mitochondrial respiration was responsible for 50% and 56% of cellular GSH in the wild type and CNTF-treated rds retinas, respectively (Fig. 7m).

**Figure 7.**
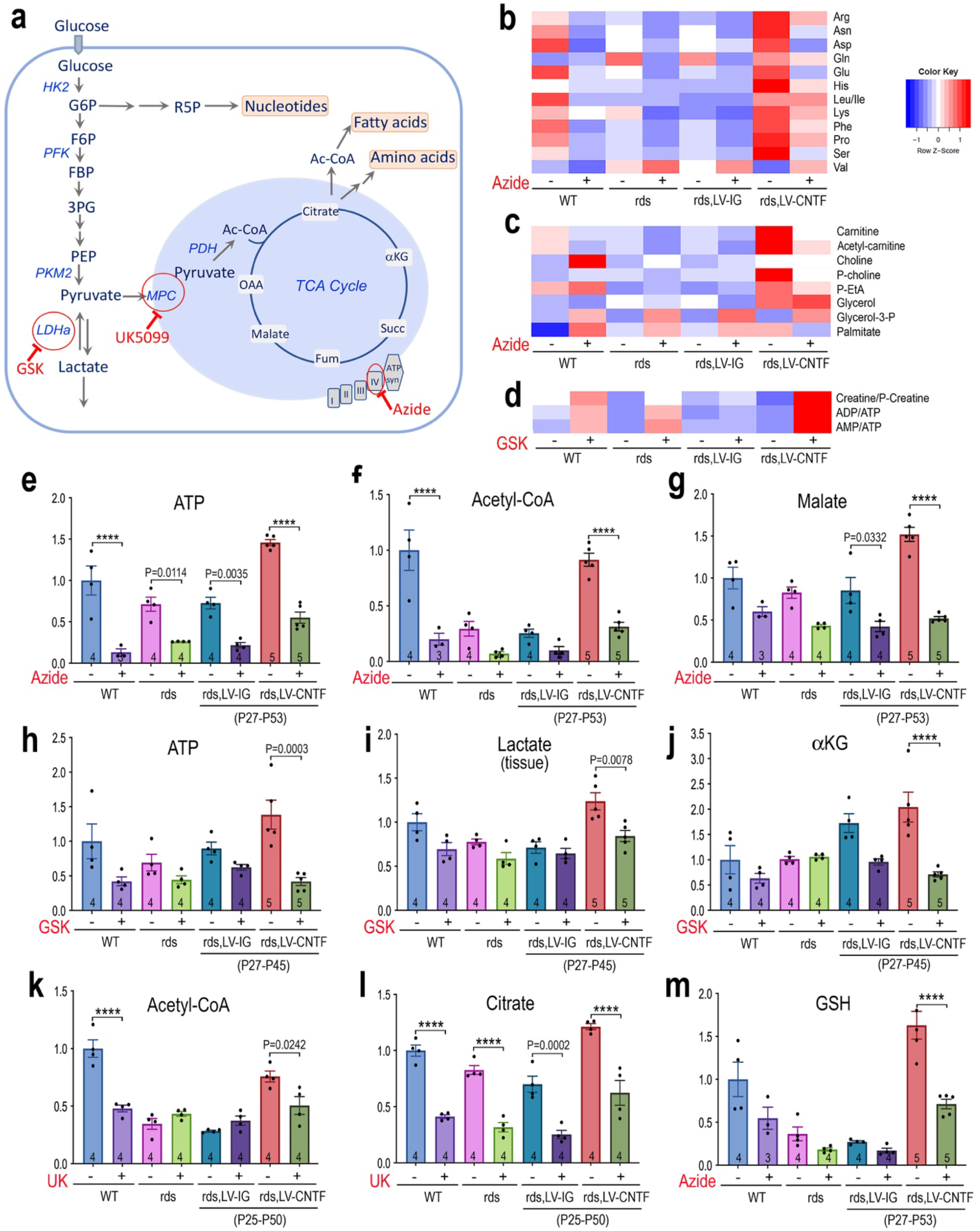
Contribution of glycolysis and mitochondrial activities to retinal metabolism and CNTF-induced metabolic changes. **a** Schematic illustration of the glycolytic pathway and TCA cycle with glucose as a fuel. Enzymatic steps affected by specific inhibitors are indicated (red circle). **b-d** Metabolomics heatmaps show effects of inhibitors in WT, rds, and rds retinas treated with LV-IG or LV-CNTF on metabolites. **b** amino acids. **c** fatty acid synthesis intermediates. **d** energy currency related metabolites. **e-m** Bar graphs show effects of inhibitors on the levels of ATP (**e, h)**, lactate in the tissue (**i**), acetyl-CoA (**f, k**), TCA cycle metabolites (**g, j, l**), and GSH (**m**). Viral vector treatment periods are shown below treated rds samples. For **e**-**m**, data are presented as mean ± SEM. Independent sample numbers (N) are indicated within the bars. Two-way ANOVA and Tukey’s multiple comparison test were applied for the entire group. For clarity, only significant adjusted P values for pairs of samples with and without a given inhibitor are shown with P < 0.0001 indicated as ****. See Source Data for the entire statistical analysis.

The glycolytic pathway enzyme lactate dehydrogenases (LDH) catalyze the conversion between pyruvate and lactate. Inhibition of LDH by GSK2837808A in the wild type retina reduced ATP level by 58%, whereas CNTF-treated rds retinas sustained 65% ATP reduction (Fig. 7h), indicating that aerobic glycolysis contributed to a larger portion of ATP in CNTF-treated rds retinas. Inhibiting LDH not only increased the AMP to ATP ratios, but also led to increased creatine to p-creatine ratios (Fig. 7d). Furthermore, metabolomics analysis showed significant increases of lactate in CNTF-treated rds retinas compared to rds or rds treated with LV-IG (Fig. 7i), indicating an enhanced glycolysis. Consistent with the conversion from lactate into pyruvate to supply mitochondria, inhibiting LDH activity also led to a significant reduction of the TCA cycle product alpha ketoglutarate (Fig. 7j), likely due to a hindered pyruvate production.

To further examine pyruvate utilization in mitochondrial metabolic processes, we applied the mitochondrial pyruvate carrier inhibitor UK5099, which caused a reduction of acetyl-CoA (Fig. 7k) and TCA cycle intermediate citrate (Fig. 7l), confirming that pyruvate transport to mitochondria supported more than 50% of the citrate production in both the wild type and rds mutant.

### Utilization of fatty acid oxidation by wild type and degenerating retinas

Since the neural retina has highly active lipid biosynthesis and turnover [73], we examined the wild type and rds mutant retinas in their abilities to utilize palmitate as a fuel source in the absence of glucose in the medium (Fig. 8a; Source Data for Fig. 8). CNTF-treated rds retinas showed higher than wild type levels of amino acids using palmitate (Fig. 8b). Inhibiting the carnitine palmitoyltransferase 1 (CPT1) with etomoxir resulted in marked reduction of amino acids in both wild type and CNTF-treated rds retinas (Fig. 8b), revealing that the retina relied substantially on fatty acid oxidation under the glucose deprivation condition. As expected, CPT inhibition led to increase levels of carnitine and reduced levels of acetyl carnitine in both wild type and CNTF-treated rds retinas, (Fig. 8c, 8i), reflecting disruption of fatty acid transport from the cytoplasm into mitochondria. Compared with the wild type, the rds retina exhibited a nearly 2-fold capacity to produce ATP when palmitate was supplied (Fig. 8d). CNTF-treated rds retina retained the efficiency to synthesize ATP using palmitate, but sustained more severe deficits with etomoxir treatment (Fig. 8d). CPT1 inhibition also nearly demolished p-creatine generation in the wild type and CNTF-treated rds retinas (Fig. 8e), confirming the potential contribution of fatty acid beta-oxidation to the retinal energy buffer. CPT1 inhibition did not affect glycolytic pathway metabolites such as 3PG (Fig. 8f), but did reduced the level of citrate, which can be driven by fatty acid beta-oxidation (Fig. 8g).

**Figure 8.**
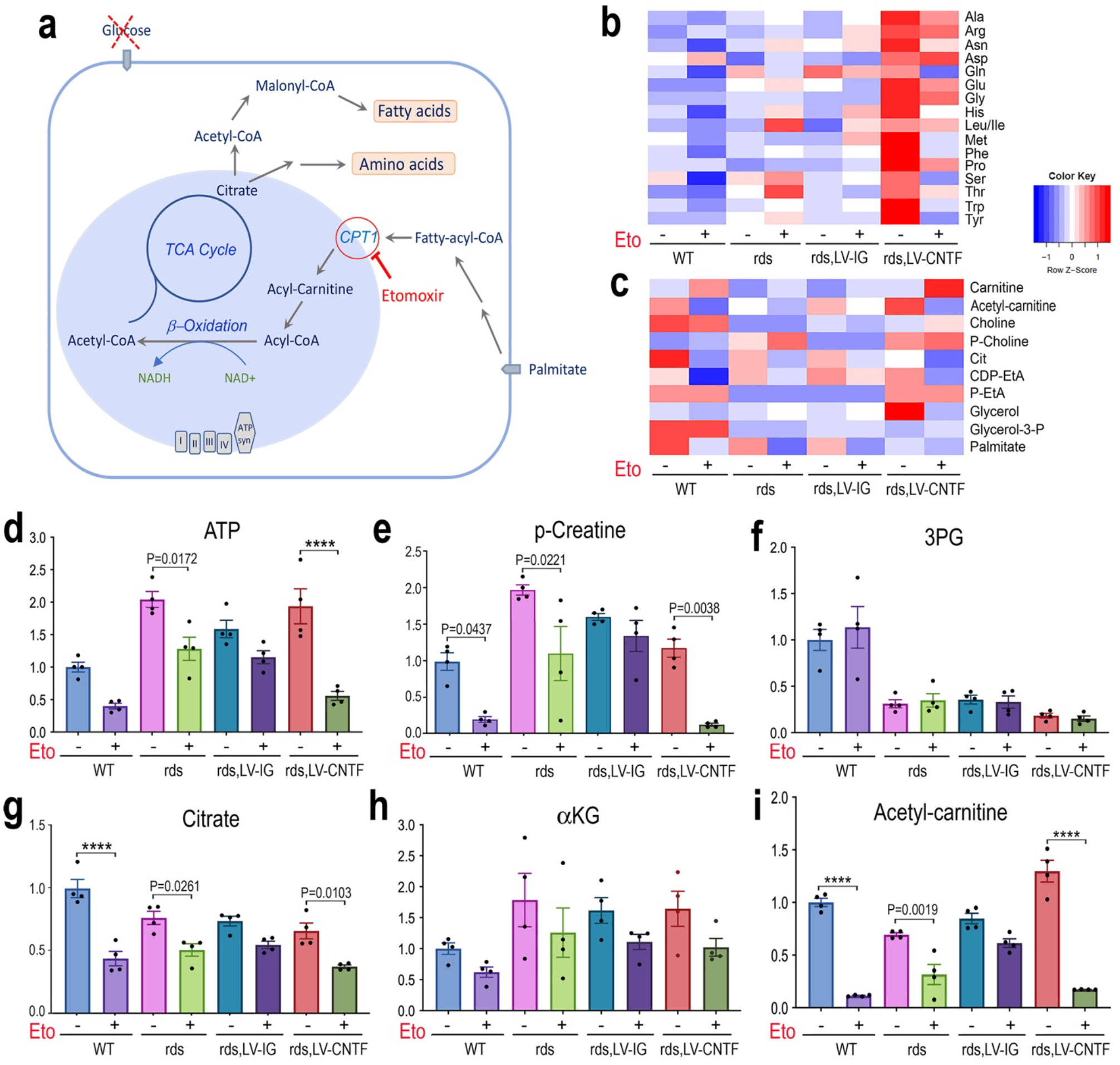
Consumption of fatty acid fuel by wild type and rds mutant retinas. **a** Schematic illustration of fatty acid metabolic pathways and Cpt1 as a target of inhibitor Etomoxir. **b, c** Metabolomics heatmaps show effects of Etomoxir on levels of (**b**) amino acids and (**c**) fatty acid synthesis intermediates in WT, rds, and rds retinas treated with LV-IG or LV-CNTF from P28-P53. **d-i** Bar graphs show effects of Etomoxir on levels of ATP (**d**), p-creatine (**e**), 3PG (**f**), citrate (**g**), αKG (**h**) and acetyl-carnitine (**i**). Independent samples N=4 for all conditions. For **d-i,** data are presented as mean ± SEM. Two-way ANOVA and Tukey’s multiple comparison test were applied for the entire group. For clarity, only significant adjusted P values for pairs of samples with and without Etomoxir are shown with P < 0.0001 indicated as ****. See Source Data for the entire statistical analysis.

## DISCUSSION

In this study, we provide evidence that delivery of the same CNTF used in human trials to a mouse model of retinitis pigmentosa results in a significant change in retinal metabolism, thus revealing a previously unknown cellular mechanism underlying the potent and broad neurotrophic effects of CNTF.

Cellular respiration assays using retinal tissues detected early and persistent decreases in oxygen consumption and elevation of extracellular acidification in the rds mutant. These changes likely reflect the adaptation of the retinal metabolism under the condition of progressive photoreceptor loss. SIM imaging revealed fragmental and ectopically distributed rod mitochondria, suggesting a possible decline of mitochondrial functionality in the rds mutant. Both SIM and TEM analyses showed that CNTF treatment partially restored the morphology of mitochondria in the rod inner segment, the major cellular biosynthetic site for photoreceptors. However, instead of improving mitochondrial respiration, our results indicated that CNTF treatment caused a reduction of ATP-linked OCR. Furthermore, direct measurements of mitochondrial respiratory chain activities confirmed the suppression of complex I and complex IV functions following CNTF treatment. Concomitant with the suppression of respiratory chain activity, we detected significantly increased ECAR following CNTF treatment, suggesting the likelihood of enhanced glycolysis.

Our biochemical and molecular analyses indeed support the above conclusion and provide a comprehensive view of the retinal metabolic status in the rds mutant and under the influence of CNTF. Metabolomics analyses using glucose as a fuel revealed deficiencies in ATP, amino acids, and fatty acid biosynthetic intermediates, as well as a severe reduction in the major antioxidant glutathione in the rds mutant retina. CNTF treatment effectively improved the retinal energy supply resulting in more than a 50% increase of ATP and a three-fold increase of p-creatine to the wild type levels. In addition, we detected significant elevation of glycolytic pathway intermediates, TCA cycle metabolites, as well as various anabolic metabolites, including amino acids, lipid biosynthetic intermediates, and nucleotide derivatives. Strikingly, CNTF treatment also restored the important antioxidant glutathione to the level found in wild type retinas. This is not surprising as glutathione biosynthesis requires amino acids glutamate, glycine, and cysteine [74]. Consistent with the metabolomics data, molecular analyses also revealed increased expression of glycolytic pathway and TCA cycle gene transcripts, and active forms of glycolytic pathway enzymes. Taken together, these results provide strong evidence that CNTF signaling in the degenerating retina leads to a global metabolic alteration by promoting aerobic glycolysis and anabolism. Since metabolomics and transcriptome analyses performed used the entire retina, the results described reflect the summation of responses by various retinal cell types to CNTF rather than photoreceptors alone. Future studies combining genetics and multiomics approaches are necessary to determine cell type-specific responses to exogenous CNTF.

Metabolomics analyses using glucose in conjunction with metabolic inhibitors permitted us to determine the contributions of glycolysis and mitochondrial respiration to the energy supply of healthy and degenerating retinas. Our results showed a reduced reliance on mitochondrial respiration in CNTF-treated rds retina to generate high levels of ATP and p-creatine, thus further supporting the role of CNTF-dependent elevation of aerobic glycolysis. Consistent with the elevated ECAR in CNTF-treated rds mutant, metabolomics analysis detected elevated lactate in retinal tissues. Further, LDHa transcripts and the active form of LDHa are both increased under CNTF influence. Since LDH catalyzes a bidirectional reaction, it is conceivable that CNTF may have promoted the conversion of lactate toward pyruvate. Notably, the lactate to pyruvate enzymatic reaction also produces NADH for cellular metabolism and as an electron donor in oxidative phosphorylation. Indeed, inhibition of the LDH enzymatic activity resulted in a 58% reduction of ATP and 65% decrease of alpha-ketoglutarate, indicating that LDH-catalyzed pyruvate production was critical for CNTF-dependent energy production as well as the mitochondrial TCA cycle. Consistently, inhibiting the mitochondrial pyruvate carrier led to significant reduction of acetyl-CoA and citrate. Moreover, in the presence of the complex IV inhibitor azide, we detected not only reduced TCA cycle products, but also amino acids and lipid biosynthetic intermediates, as the production of these anabolic metabolites relies on the supply of acetyl-CoA and the TCA cycle metabolites. These findings demonstrate that CNTF not only upregulates aerobic glycolysis and dampens mitochondrial respiratory chain activity, but also strongly impacts TCA cycle activities that are critical for amino acid and lipid biosynthesis.

It is known that in the case of retinitis pigmentosa, rod photoreceptor death leads to cone cell starvation of glucose [54, 55]. In the absence of glucose supply, we found that the rds retina displayed an increased capacity to utilize palmitate to produce ATP and p-creatine compared to the wild type. Blocking palmitate transport into the mitochondria resulted in decreases of ATP and p-creatine production in both wild type and CNTF-treated rds retinas. Interestingly, when provided with palmitate, CNTF-treated rds retinas showed only increased amino acid contents without elevations of other anabolic metabolites compared to using glucose as a fuel. Consistent with retinal consumption of fatty acids through beta-oxidation and the TCA cycle, inhibition of palmitate transport also caused the reduction of citrate, amino acids, and most lipid biosynthetic pathway intermediates, but had no effect on the levels of glycolytic pathway metabolites. Therefore, compared to the wild type, the rds mutant retina can more efficiently use fatty acids as a fuel source when experiencing glucose deprivation.

The CNTF-triggered retinal metabolic changes show striking similarities to the metabolic signatures of cancer cells as described by Warburg nearly a century ago [75]. We have demonstrated previously that CNTF primarily activates the Jak-STAT and ERK pathways in the retina [38–40]. Activation of STAT3 has long been deployed to maintain the pluripotency of stem cells, which rely heavily on aerobic glycolysis to supply energy and various metabolites for proliferation, epigenetic modification, and cellular anabolic processes [76, 77]. Elevated STAT3 signaling has been associated with the oncogenic process [78–80]. Intriguingly, in addition to the nuclear localized pY705 STAT3 dimer to regulate target gene transcription, the Ser727 phosphorylated STAT3 has been shown to enter mitochondria and enhance electron transport chain function, as well as promote oncogenic transformation [81, 82]. However, the exact role of pS727 STAT3 in mitochondrial function remains to be resolved [83–85]. A previous study has shown that elevating wild type or a constitutively active STAT3 in mutant mouse rod photoreceptors can delay degeneration, and that the pY705 STAT3 mediates this protective effect [86]. Since CNTF signaling asserts a strong influence on the retinal transcriptome [44], the precise functions of pY705 and pS727 STAT3 in CNTF-mediated metabolic modulation require further investigations. Consistent with our results that exogenous CNTF suppresses mitochondrial respiratory chain activity and promotes cell viability, accumulating evidence indicates that partial uncoupling of mitochondrial respiration can reduce oxidative stress and support neuronal survival under stressful conditions [87–91]. Furthermore, blocking mitochondrial transport of pyruvate has been shown to increase glycolysis and potentiate endogenous stem cell activities [92], as well as attenuate neuronal loss [93]. It may appear paradoxical that CNTF improves mitochondrial morphology on the one hand, yet suppresses respiratory chain function. However, it is plausible that CNTF signaling results in a partial uncoupling, but does not compromise the TCA cycle activity in the mitochondrial matrix thus and enhancing anabolism.

Results of this study provide much needed insight at the molecular and biochemical levels for the ongoing clinical trials aimed at treating blinding diseases. Recent studies have shown that MacTel type 2, which causes late-onset retinal degeneration, is associated with metabolic dysfunction in the serine-glycine biosynthetic pathway [94, 95]. In addition, metabolic changes in phospholipids including phosphatidylethanolamines have been detected among MacTel patients [96]. Based on our findings, one possible explanation for the observed efficacy of the phase II CNTF trial for MacTel type 2 [31] is likely due to CNTF-dependent alterations of retinal metabolism, especially the elevation of retinal amino acid contents. CNTF is known to have neurotrophic effects on RGC survival and RGC axon regeneration [13, 19]. Future investigations examining retinal cell type specific effects of CNTF will facilitate elucidating cellular mechanisms for the on-going CNTF glaucoma trials. In summary, findings of this study have revealed cellular responses of the neural retina to CNTF treatments, thus suggesting potential therapeutic treatments for neurodegenerative diseases.

## METHODS

### Animals

Mice were kept on a 12/12-hour light/dark cycle with Rodent Diet 20 (Pico-Lab, catalog number 15053). The Rds transgenic mice carrying the Prph2(P216L) mutation [64] were maintained by crossing with the wild type CD1 mice from the Jackson Laboratory. Progeny of the cross without the Prph2(P216L) transgene were used as wild type littermate controls. Mice expressing rod photoreceptor specific mitochondrial PhAM reporter were generated by crossing mice that carry both Rho iCre75 [65] (Jackson Laboratory Stock No.015850) and the Rds/Prph2(P216L) transgenes with the PhAM reporter mice [66] (Jackson Laboratory Stock No.018385). Genotyping was performed by tail genomic DNA extraction followed by PCRs using primers listed in Supplementary Table 1a. Equal numbers of male and female mice are used for a given assay. All animal procedures followed National Institutes of Health guidelines and were approved by the Animal Research Committee at University of California Los Angeles under the protocol number ARC-1996-047.

### Lentiviral vector production and intraocular delivery

The lentivirus LV-CNTF and the control lentivirus LV-IG share the same vector backbone and encode the CMV promoter followed by CNTF-IRES-GFP or IRES-GFP, respectively [40]. The LV-CNTF expresses the same secreted form of human CNTF with S166D and G167H substitutions used in clinical trials [42]. The lentiviral vectors and helper plasmids were used to co-transfect HEK293t cells, and the media were collected 48 hour post-transfection. Ultracentrifugation-concentrated viral particles were resuspended and used to obtain viral titers by serial dilution and infection of HEK293t cells followed by immunolabeling and quantification of GFP-positive clones [40, 97]. For in vivo delivery, lentiviral stocks with titers of 1 × 10^7^ CFU/ml were injected subretinally at 0.5 μl per eye.

### Confocal, super-resolution, and transmission electron microscopies

Cells or retinal tissues were fixed with 4% (wt/vol) paraformaldehyde in PBS and processed as described previously [39]. Confocal fluorescent images were captured using Olympus FluoView 1000 scanning laser confocal microscope. Super-resolution images were captured with either Zeiss Airyscan LSM 800 or General Electric DeltaVision OMX microscope for structure illumination microscopy (SIM) using PlanApoN 60x/1.42 NA oil objective (Olympus). For Airyscan imaging, retina was dissociated using papain as described [98]. For SIM imaging, 14 μm thickness cryosections were adhered to #1.5 coverslip (ThermoFisher Scientific) coated with Matrigel (BD Biosciences and StemCell Technologies) and mounted on glass slide using Vectashield mounting medium (Vectorlabs). The coverslip/slide was sealed with CoverGrip (Biotium). Images were acquired in 3D-SIM mode using a Z-spacing of 0.125 μm, and reconstructed using Softworx software (GE Healthcare). Imaris software (Oxford Instruments) was used to extract 5 μm thickness 3D images and create 3D rotating video clips. For TEM, eyes were fixed in 2% (wt/vol) formaldehyde and 2.5% (wt/vol) glutaraldehyde in 0.1 M sodium phosphate buffer and processed as described [99]. TEM images were acquired using JEM-1400 (JEOL) electron microscope.

### Mitochondrial morphology analysis

SIM images were analyzed in FIJI/ImageJ [100] using the open-source software plugin MitoMap (http://www.gurdon.cam.ac.uk/stafflinks/downloadspublic/imaging-plugins)[68]. The plugin automates the process to compute mitochondrial volume (μm^3^), surface area (μm^2^), and other geometrical features in a region of interest (ROI). For each sample analyzed, the 32-bit OMX images stack (dv file) are loaded onto ImageJ using the Bio-formats plugin [101], and an ROI was chosen that includes the photoreceptor inner segment (40 × 15 × 10 μm^3^) region. The MitoMap plugins converted the images stack to 16-bit and applied Otsu thresholding [102] to extract the PhAM reporter-labeled mitochondrial volume. Before the quantification analysis, objects with volume smaller than 0.1 μm^3^ were excluded to eliminate artifacts. The principal component analyses were performed on R Studio using ‘factoextra’ and ‘FactoMineR’ packages [103].

### Seahorse assays

For tissue OCR and ECAR assays, retinas were dissected in HBSS and tissue disks containing all retinal layers were obtained using 1 mm diameter biopsy puncher (Miltex) [104]. Retinal disks were washed with unbuffered Seahorse XF Base DMEM pH 7.4 (Agilent) and placed in a well of 96-well microplate containing 175 μl medium supplemented with 6 mM glucose (pH 7.4). Retinal disks were incubated for 30 minutes at 37°C prior to running the assay using a Seahorse XF96 Analyzer (Agilent). During the assay, 3 μM oligomycin (Port A), 0.6 μM and 1.1 μM FCCP (Ports B and C), and 2 μM antimycin A and 2 μM rotenone (Port D) were injected into the assay medium (Supplementary Table 1b).

For isolated mitochondria seahorse assays, eyes were dissected in cold PBS and three retinas were pooled for each sample. After washing and removal of PBS, 500 μl of MSHE buffer (210 mM mannitol, 70 mM sucrose, 5 mM HEPES, 1 mM EDTA, 0.5% BSA, pH 7.2) was added to the pooled retinas. Retinal tissues were homogenized by drawing through a 23G needle 20 times, followed by centrifugation at 800 g for 10 minutes at 4°C. The supernatants containing mitochondria were collected and transferred to a new tube, and the pellets were homogenized and centrifuged again to re-collect the supernatants. The supernatants were then combined and centrifuged at 8000 g for 10 minutes at 4°C. The resulting pellets containing mitochondria were resuspended in 800 μl of MSHE and centrifuged again at 8000 g for 10 min at 4°C. After discarding the supernatant, the mitochondria pellet was resuspended in 35 μl of mitochondrial assay solution (MAS; 220 mM mannitol, 70 mM sucrose, 10 mM potassium phosphate monobasic, 5 mM magnesium chloride, 2 mM HEPES, 1 mM EGTA, pH7.2) without BSA. Mitochondrial protein contents were measured using the BCA assay (Pierce). Isolated mitochondria were loaded into an ice-cold Seahorse XF96 microplate with 15 μl of 10x substrate solution (50 mM pyruvate and 50 mM malate, or 50 mM succinate and 20 μM rotenone). For Complex I-driven respiration (pyruvate + malate), 7 μg protein was loaded per well; while 4 μg protein/well was loaded for Complex II-driven respiration (succinate + rotenone). The volume was adjusted to 20 μl per well and the mitochondria plate was centrifuged at 2100 g for 5 minutes to allow mitochondria to adhere to the bottom of the well. After centrifugation, the total volume of the well was adjusted to 130 μl with ice-cold MAS with 0.1% fatty acid-free BSA (pH 7.2 adjusted with 1 M KOH). During the assay, compounds were injected from the ports of the XF96 Analyzer (Source Data Table S8.B). The isolated mitochondria conditions include injection of: 1) 4 mM ADP in the presence of pyruvate and malate or succinate and rotenone (State 3: maximal ATP synthesis capacity), 2) 3μM of the ATP Synthase inhibitor, oligomycin (State 4o: proton leak), 3) the chemical uncoupler, FCCP at 4 μM, and 4) 2μM of the Complex III inhibitor, antimycin. When measuring Complex IV respiration, isolated mitochondria conditions include injection of: 1) 4 mM ADP in the presence of pyruvate and malate or succinate and rotenone (State 3: maximal ATP synthesis capacity), 2), 3) 1 mM TMPD to (donate electrons to cytochrome c/Complex IV) with ascorbate to keep TMPD in the reduced state, and 4) 40 mM of the Complex IV inhibitor, azide.

### Metabolomics analysis

For polar metabolite extraction from retinal tissue and medium, retinas were dissected in cold Krebs-Ringer Bicarbonate Buffer pH 7.4 (KRB: 119.78 mM NaCl, 2.6 mM CaCl_2_, 4.56 mM KCl, 0.49 mM MgCl_2_, 0.7 mM Na2HPO_4_, 1.3 mM NaH_2_PO_4_, 14.99 mM NaHCO_3_ and 10 mM HEPES) with 5 mM substrates (glucose or palmitate). Whole retina or 1 mm retinal disks were incubated in 500 μl of KRB with or without inhibitors (Supplementary Table 1c), for 60 minutes at 37°C. For metabolites in the medium, 20 μl of medium was collected and mixed with 500 μl of cold 80% methanol by vortex. For tissue metabolites, after removal of the KRB medium, retinal tissues were homogenized in 500 μl of cold 80% methanol by drawing through 23G needle 20 times. Both the medium and tissue extracts were spun in a microfuge at 13,000 rpm at 4°C for 10 minutes. The supernatants were transferred into glass vials, 2 nmoles of norvaline was added to each vial, and the extracts were desiccated using EZ-2Elite evaporator and stored at −80°C.

Dried metabolites were resuspended in 50% ACN:water and 1/10^th^ was loaded onto a Luna 3 μm NH2 100A (150 × 2.0 mm) column (Phenomenex). The chromatographic separation under HILIC conditions was performed on a Vanquish Flex (Thermo Scientific) with mobile phases A (5 mM NH4AcO pH 9.9) and B (ACN) and a flow rate of 200 μl/min. A linear gradient from 15% A to 95% A over 18 min was followed by 9 min isocratic flow at 95% A and re-equilibration to 15% A. Metabolites were detection with a Thermo Scientific Q Exactive mass spectrometer run with polarity switching (+3.5 kV/− 3.5 kV) in full scan mode with an m/z range of 65-975. TraceFinder 4.1 (Thermo Scientific) was used to quantify the targeted metabolites by area under the curve using expected retention time and accurate mass measurements (< 5 ppm). Data analysis was performed using in-house R scripts (https://github.com/graeberlab-ucla/MetabR).

### RNA sequencing

The whole retinal transcriptome analysis was performed as described [44]. Briefly, total RNA was isolated from wild type, and rds/Prph2(P216L) mutant retinas treated with LV-IG or LV-CNTF from P25-P35 (N = 3, each N contains 2 retinas). Sequencing libraries were prepared using the Illumina TruSeq Stranded Total RNA with Ribo-Zero Gold Library Prep kit. Paired-end sequencing of 69 bp of the libraries was performed using HiSeq 4000 Sequencer (Illumina). After quality control and filtering, the median size was 5.65 Gb per library (range 3.66–7.26 Gb). Sequenced reads were then aligned to the Mouse reference mm10 from UCSC (genome-euro.ucsc.edu) using Hisat2 (v2.0.4) [105]. The expression levels were normalized by calculating the fragments per kilobase million reads (FPKM) values.

### Western blots

For western blot analysis, dissected retina was washed with PBS, then lysed with RIPA (50 mM Tris, 150 mM NaCl, 1% Triton X-100, 2% BSA and complete protease inhibitor cocktail, pH 7.4). The lysates were then resolved on SDS-PAGE and western blots were performed as described [40]. The protein signals were imaged using Odyssey^®^ CLx Imaging System (LI-COR Biosciences). Primary and secondary antibodies used are summarized in Supplementary Table.

### Statistics

All N numbers represent independent samples analyzed, as indicated pertinently in each figure, including Supplementary Figures. All error bars in bar graphs of figures and supplementary figures are presented as mean value ± SEM. For Figure 3a, a two-tailed student’s t-test was used to compare the wild type with rds mutant data (Source Data for Fig. 3). For the rest of the data in various figures, one-way or two-way ANOVA and Tukey’s multiple comparison tests, when appropriate, were performed (see Source Data for detailed statistics for each figure). Actual P values are shown in figures with P < 0.05 considered statistically significant and P<0.0001 indicated as ****.

## DATA AVAILABILITY

RNA-sequencing data have been deposited at Gene Expression Omnibus (GEO) and will be publicly available with Accession numbers GSE216208. Data user can retrieve raw sequence data at https://urldefense.com/v3/_https://www.ncbi.nlm.nih.gov/geo/query/acc.cgi?acc=GSE216208_;!!F9wkZZsI-LA!G70HdsG0Nn3a6tDxjMuIflgFcvKAk-XtN9FAC_B_H2tkiMCE8s-u5JxfL-HvFIA7sMyEXhIBk9jKX3xl-iYC$ [ncbi[.]nlm[.]nih[.]gov]

All data generated for this study are included in the main and supplementary figures. For all quantitative figures, data of individual values as well as the results of statistical tests are provided in the Source Data files with the paper. Other data that support the findings of this study are available on request from the corresponding author.

## CODE AVAILABILITY

Images analysis of mitochondria used FIJI/ImageJ [97] open-source software plugin MitoMap (http://www.gurdon.cam.ac.uk/stafflinks/downloadspublic/imaging-plugins)[68].

Code used for metabolomics data analysis is deposited in Github Repository: https://github.com/graeberlab-ucla/MetabR.

## ACKNOWLEDGMENTS

This work was supported by NIH grant R01EY026319 to X.J.Y., NIH core grant P30EY000331, and an unrestricted grant from the Research to Prevent Blindness to the Department of Ophthalmology at University of California Los Angeles.

## AUTHOR CONTRIBUTIONS

K.D.R and X.J.Y., conception and research designs; K.D.R and T.T.T.N., SIM and MitoMap analysis; D.B., electron microscopy; K.D.R. and Y.W., transcriptome analysis; L.S., L.V., mitochondrial respiration; K.R., O.S., data interpretation; K.D.R. and X.Z., Western blot analysis; J.t.H., metabolomics; K.D.R. and X.J.Y., data analysis and figure preparation; X.J.Y., manuscript writing.

---

**Supplementary Figure 1.**
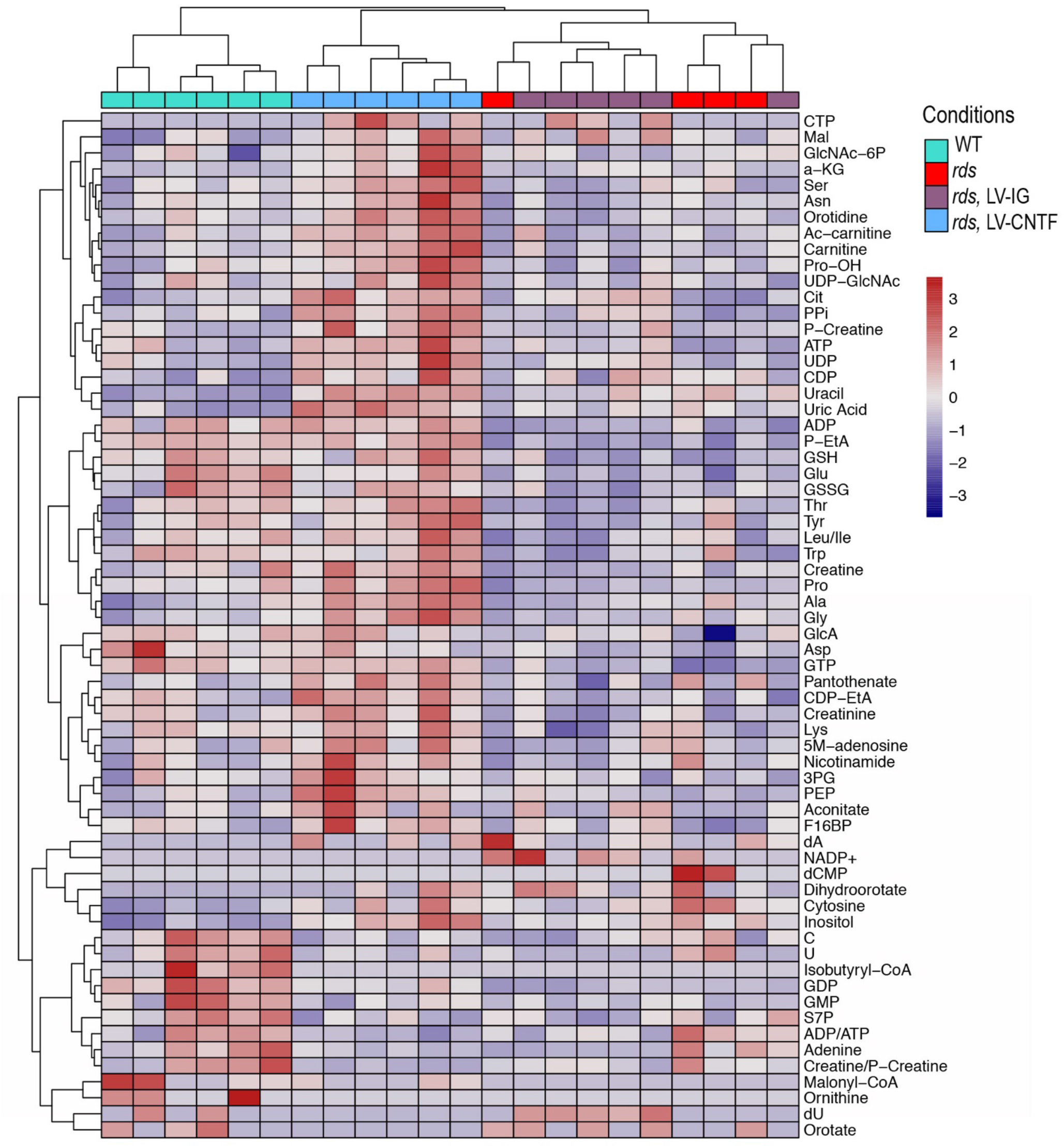
Metabolomics heatmap of WT, rds, and rds retinas treated with LV-IG or LV-CNTF. Rds retinas were treated with LV-IG or LV-CNTF from P25 to P36. Retinal tissues were incubated with 5mM glucose for 60 minutes prior to metabolomics analysis. Independent retina samples: N=6 for WT; N=4 for rds; N=6 for rds treated with LV-IG or LV-CNTF.

**Supplementary Figure 2.**
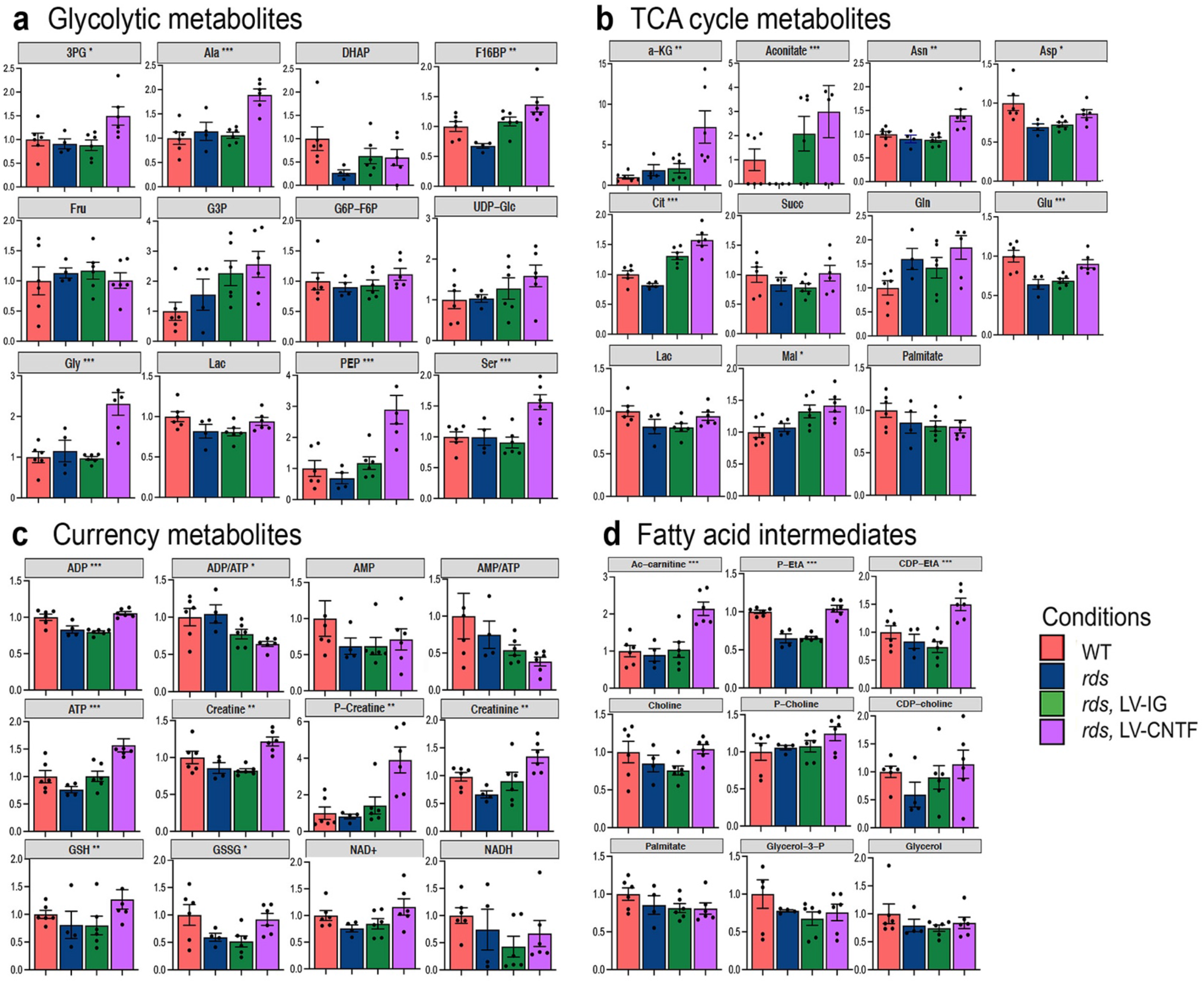
Bar graphs of glycolysis, TCA cycle, currency, and fatty acid metabolites. Rds retinas were treated with LV-IG or LV-CNTF from P25 to P36. Retinal tissues were incubated with 5mM glucose for 60 minutes prior to metabolomics analysis. Independent retina samples: N=6 for WT; N=4 for rds; N=6 for rds treated with LV-IG or LV-CNTF. For **a-d**, data are presented as mean ± SEM. One-way ANOVA was used to detected statistical significance in each metabolite group. * P < 0.05; ** P < 0.01; *** P< 0.001.

**Supplementary Figure 3.**
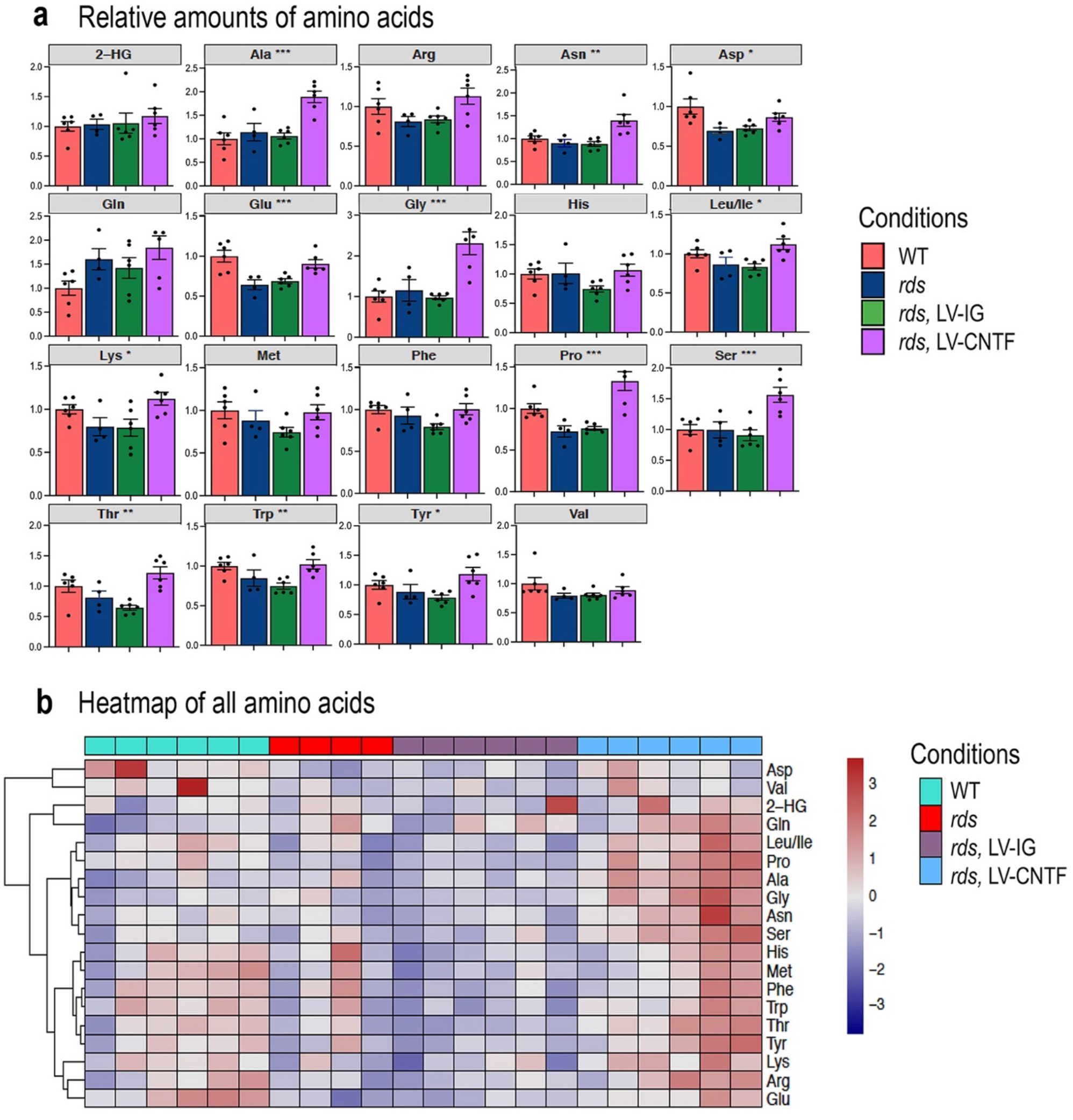
Bar graphs and heatmap of amino acids with glucose as a nutrient. Rds retinas were treated with LV-IG or LV-CNTF from P25 to P36. Retinal tissues were incubated with 5mM glucose for 60 minutes prior to metabolomics analysis. Independent retina samples: N=6 for WT; N=4 for rds; N=6 for rds treated with LV-IG or LV-CNTF. **a.** Bar graphs of retinal contents of amino acids. Data are presented as mean ± SEM. One-way ANOVA was used to detected statistical significance in each metabolite group. * P < 0.05; ** P < 0.01; *** P< 0.001. **b.** Heatmap of relative amounts of amino acids.

**Supplementary Figure 4.**
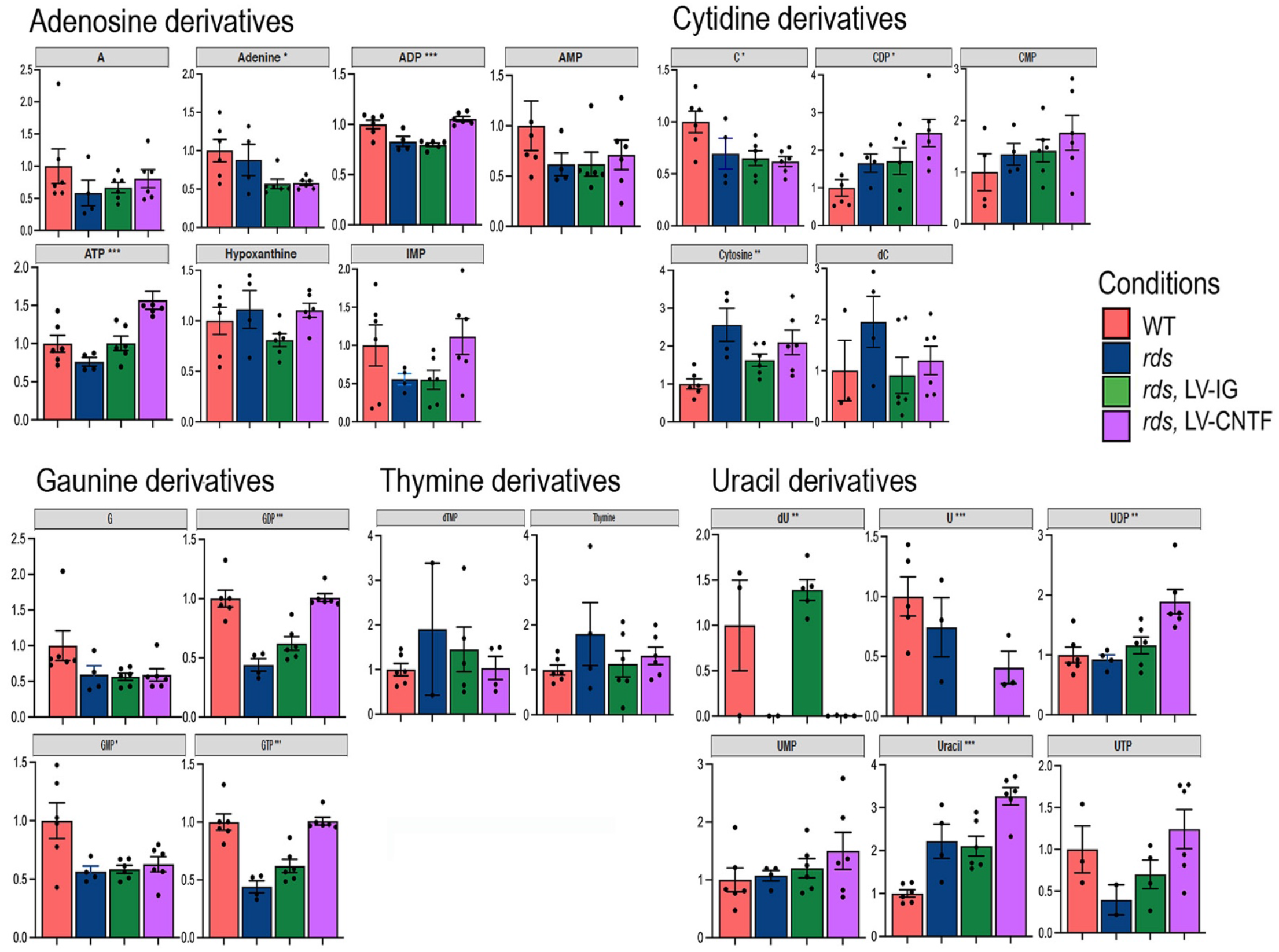
Bar graphs of nucleotides and derivatives with glucose as a nutrient. Rds retinas were treated with LV-IG or LV-CNTF from P25 to P36. Retinal tissues were incubated with 5mM glucose for 60 minutes prior to metabolomics analysis. Independent retina samples: N=6 for WT; N=4 for rds; N=6 for rds treated with LV-IG or LV-CNTF. Data are presented as mean ± SEM. One-way ANOVA was used to detected statistical significance in each metabolite group. * P < 0.05; ** P < 0.01; *** P< 0.001.

**Supplementary Figure 5.**
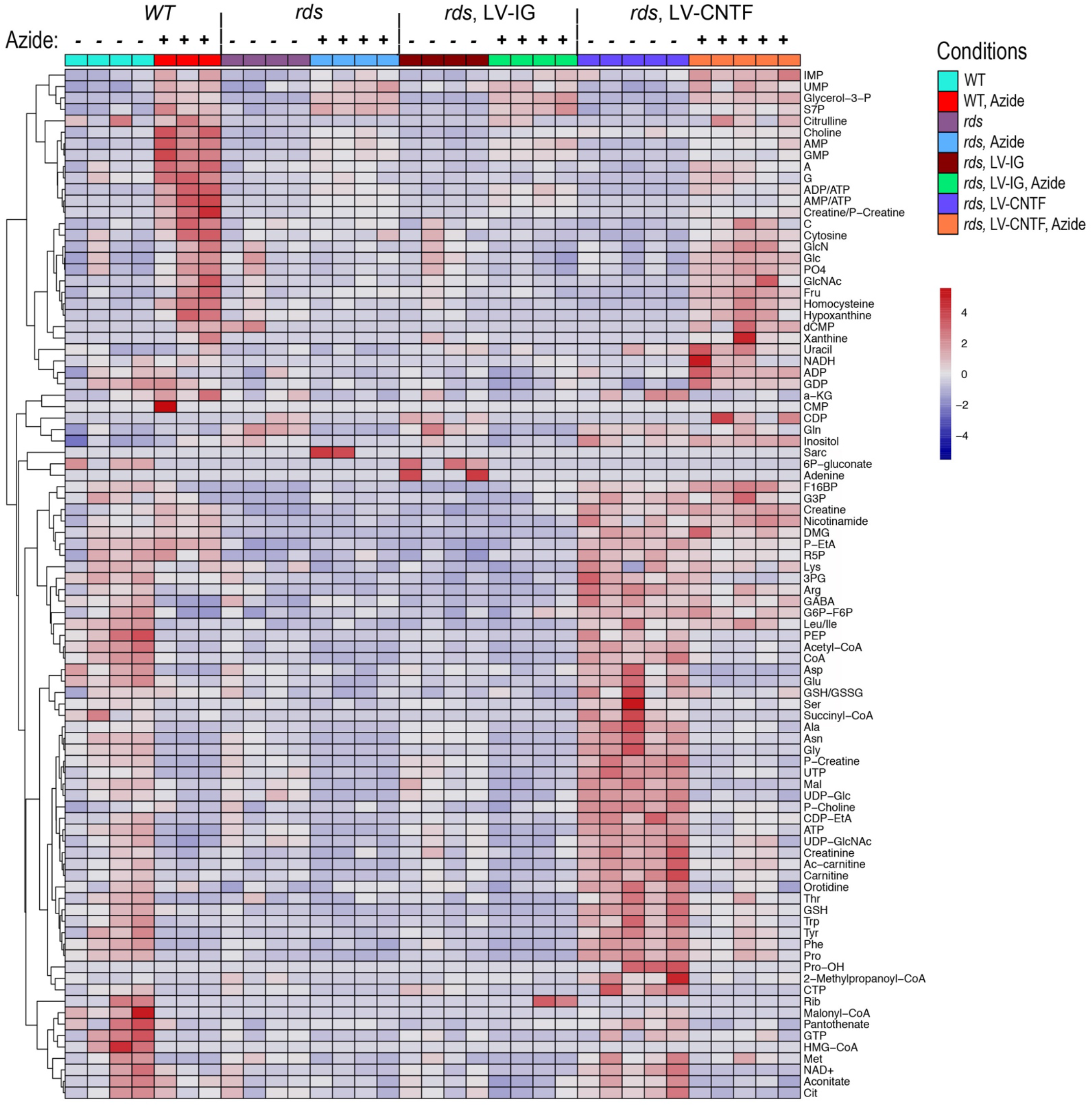
Metabolomics heatmap of WT, rds, and rds retinas treated with LV-IG or LV-CNTF and Na^+^Azide. Rds retinas were treated with LV-IG or LV-CNTF from P27 to P53. Retinal tissues were incubated with 5mM glucose for 60 minutes in the presence or absence of Na^+^Azide. Independent samples: N=3 for WT+azide; N=4 for WT-azide, rds +/−azide, rds treated with LV-IG +/−azide; N=5 for rds treated with LV-CNTF +/−azide.

**Supplementary Figure 6.**
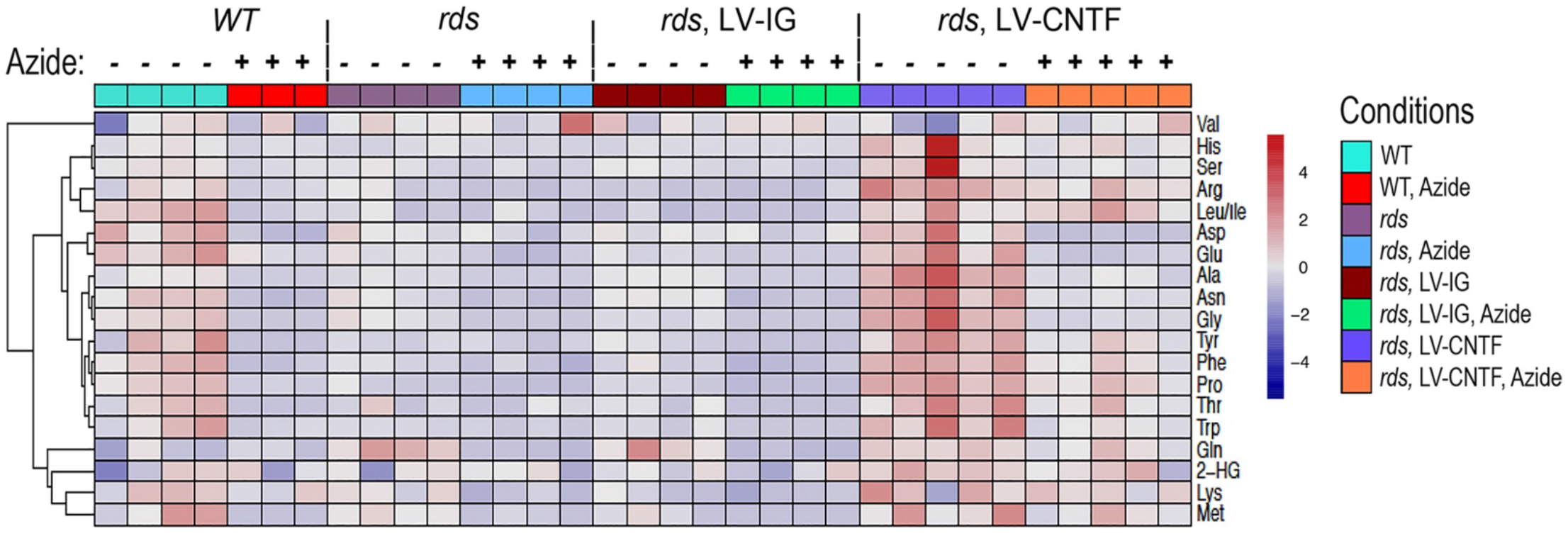
Metabolomics heatmap of amino acids with glucose and Na^+^Azide. Rds retinas were treated with LV-IG or LV-CNTF from P27 to P53. Retinal tissues were incubated with 5mM glucose for 60 minutes in the presence or absence of Na^+^Azide. Independent samples: N=3 for WT+azide; N=4 for WT-azide, rds +/−azide, rds treated with LV-IG +/−azide; N=5 for rds treated with LV-CNTF +/−azide. Relative amounts of amino acids in retinal tissues are shown as a heatmap.

**Supplementary Table.**
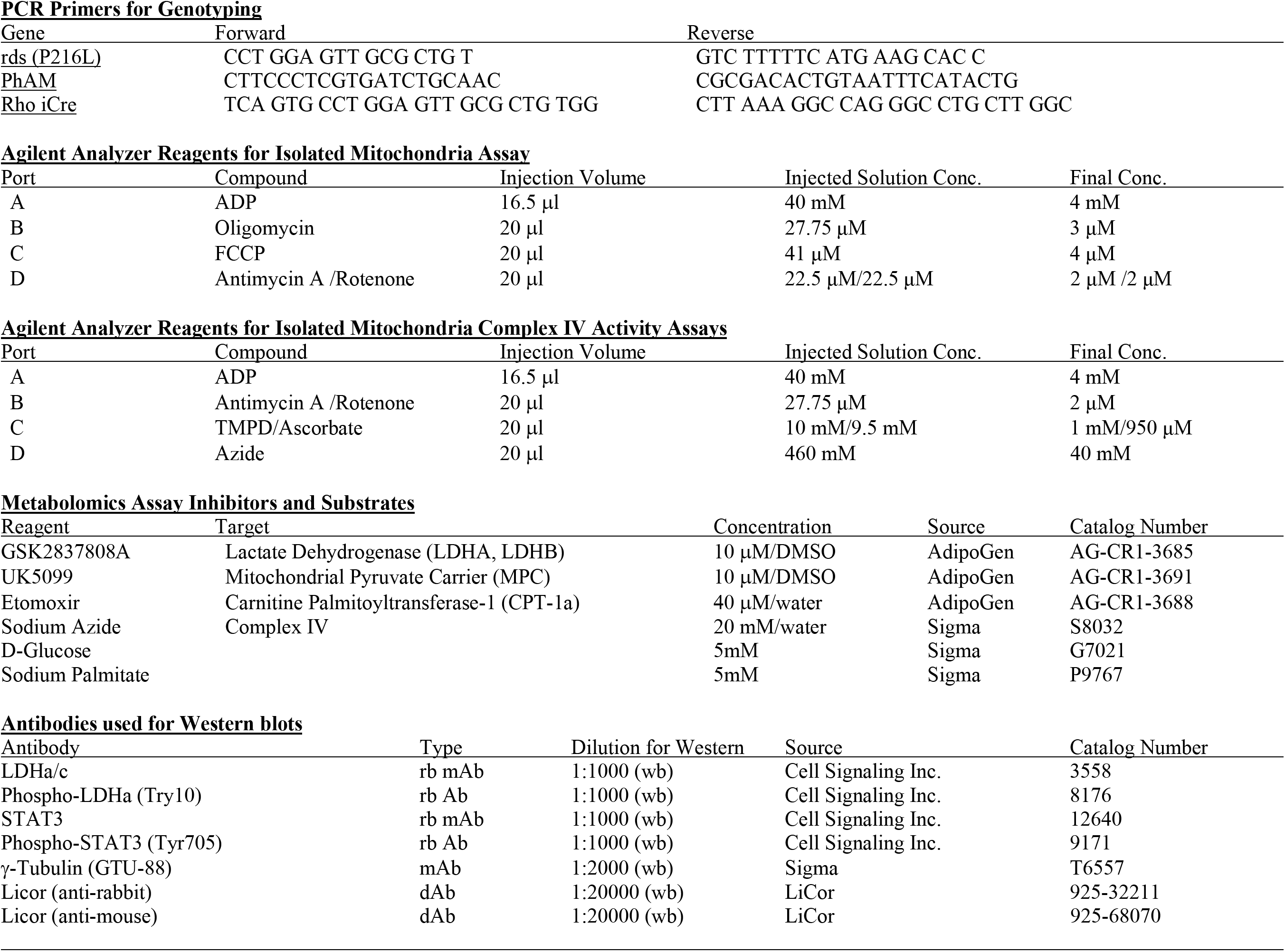
Summary of Reagents.

## REFERENCES

1. LaVail, M.M., et al., Multiple growth factors, cytokines, and neurotrophins rescue photoreceptors from the damaging effects of constant light. Proc Natl Acad Sci U S A, 1992. 89(23): p. 11249–53.

2. Wen, R., et al., CNTF and retina. Prog Retin Eye Res, 2012. 31(2): p. 136–51.

3. Cayouette, M. and C. Gravel, Adenovirus-mediated gene transfer of ciliary neurotrophic factor can prevent photoreceptor degeneration in the retinal degeneration (rd) mouse. Hum Gene Ther, 1997. 8(4): p. 423–30.

4. Adamus, G., et al., Anti-apoptotic effects of CNTF gene transfer on photoreceptor degeneration in experimental antibody-induced retinopathy. J Autoimmun, 2003. 21(2): p. 121–9.

5. Huang, S.P., et al., Intraocular gene transfer of ciliary neurotrophic factor rescues photoreceptor degeneration in RCS rats. J Biomed Sci, 2004. 11(1): p. 37–48.

6. Liang, F.Q., et al., AAV-mediated delivery of ciliary neurotrophic factor prolongs photoreceptor survival in the rhodopsin knockout mouse. Mol Ther, 2001. 3(2): p. 241–8.

7. Schlichtenbrede, F.C., et al., Intraocular gene delivery of ciliary neurotrophic factor results in significant loss of retinal function in normal mice and in the Prph2Rd2/Rd2 model of retinal degeneration. Gene Ther, 2003. 10(6): p. 523–7.

8. Bush, R.A., et al., Encapsulated cell-based intraocular delivery of ciliary neurotrophic factor in normal rabbit: dose-dependent effects on ERG and retinal histology. Invest Ophthalmol Vis Sci, 2004. 45(7): p. 2420–30.

9. Komaromy, A.M., et al., Transient photoreceptor deconstruction by CNTF enhances rAAV-mediated cone functional rescue in late stage CNGB3-achromatopsia. Mol Ther, 2013. 21(6): p. 1131–41.

10. Aslam, S.A., et al., Cone photoreceptor neuroprotection conferred by CNTF in a novel in vivo model of battlefield retinal laser injury. Invest Ophthalmol Vis Sci, 2013. 54(8): p. 5456–65.

11. Lipinski, D.M., et al., CNTF Gene Therapy Confers Lifelong Neuroprotection in a Mouse Model of Human Retinitis Pigmentosa. Mol Ther, 2015. 23(8): p. 1308–1319.

12. Cui, Q., et al., CNTF, not other trophic factors, promotes axonal regeneration of axotomized retinal ganglion cells in adult hamsters. Invest Ophthalmol Vis Sci, 1999. 40(3): p. 760–6.

13. Meyer-Franke, A., et al., Characterization of the signaling interactions that promote the survival and growth of developing retinal ganglion cells in culture. Neuron, 1995. 15(4): p. 805–19.

14. MacLaren, R.E., et al., CNTF gene transfer protects ganglion cells in rat retinae undergoing focal injury and branch vessel occlusion. Exp Eye Res, 2006. 83(5): p. 1118–27.

15. van Adel, B.A., et al., Delivery of ciliary neurotrophic factor via lentiviral-mediated transfer protects axotomized retinal ganglion cells for an extended period of time. Hum Gene Ther, 2003. 14(2): p. 103–15.

16. Cui, Q. and A.R. Harvey, CNTF promotes the regrowth of retinal ganglion cell axons into murine peripheral nerve grafts. Neuroreport, 2000. 11(18): p. 3999–4002.

17. Sun, F., et al., Sustained axon regeneration induced by co-deletion of PTEN and SOCS3. Nature, 2011. 480(7377): p. 372–5.

18. Liu, X., P.R. Williams, and Z. He, SOCS3: a common target for neuronal protection and axon regeneration after spinal cord injury. Exp Neurol, 2015. 263: p. 364–7.

19. Williams, P.R., et al., Axon Regeneration in the Mammalian Optic Nerve. Annu Rev Vis Sci, 2020. 6: p. 195–213.

20. Tao, W., et al., Encapsulated cell-based delivery of CNTF reduces photoreceptor degeneration in animal models of retinitis pigmentosa. Invest Ophthalmol Vis Sci, 2002. 43(10): p. 3292–8.

21. Sieving, P.A., et al., Ciliary neurotrophic factor (CNTF) for human retinal degeneration: phase I trial of CNTF delivered by encapsulated cell intraocular implants. Proc Natl Acad Sci U S A, 2006. 103(10): p. 3896–901.

22. Zhang, K., et al., Ciliary neurotrophic factor delivered by encapsulated cell intraocular implants for treatment of geographic atrophy in age-related macular degeneration. Proc Natl Acad Sci U S A, 2011. 108(15): p. 6241–5.

23. Talcott, K.E., et al., Longitudinal study of cone photoreceptors during retinal degeneration and in response to ciliary neurotrophic factor treatment. Invest Ophthalmol Vis Sci, 2011. 52(5): p. 2219–26.

24. Kauper, K., et al., Two-year intraocular delivery of ciliary neurotrophic factor by encapsulated cell technology implants in patients with chronic retinal degenerative diseases. Invest Ophthalmol Vis Sci, 2012. 53(12): p. 7484–91.

25. Birch, D.G., et al., Randomized trial of ciliary neurotrophic factor delivered by encapsulated cell intraocular implants for retinitis pigmentosa. Am J Ophthalmol, 2013. 156(2): p. 283–292 e1.

26. Zein, W.M., et al., CNGB3-achromatopsia clinical trial with CNTF: diminished rod pathway responses with no evidence of improvement in cone function. Invest Ophthalmol Vis Sci, 2014. 55(10): p. 6301–8.

27. Pilli, S., R.J. Zawadzki, and D.G. Telander, The dose-dependent macular thickness changes assessed by fd-oct in patients with retinitis pigmentosa treated with ciliary neurotrophic factor. Retina, 2014. 34(7): p. 1384–90.

28. Langlo, C., et al., CNGB3-Achromatopsia Clinical Trial With CNTF: Diminished Rod Pathway Responses With No Evidence of Improvement in Cone Function. Invest Ophthalmol Vis Sci, 2015. 56(3): p. 1505.

29. Birch, D.G., et al., Long-term Follow-up of Patients With Retinitis Pigmentosa Receiving Intraocular Ciliary Neurotrophic Factor Implants. Am J Ophthalmol, 2016. 170: p. 10–14.

30. Chew, E.Y., et al., Ciliary neurotrophic factor for macular telangiectasia type 2: results from a phase 1 safety trial. Am J Ophthalmol, 2015. 159(4): p. 659–666 e1.

31. Chew, E.Y., et al., Effect of Ciliary Neurotrophic Factor on Retinal Neurodegeneration in Patients with Macular Telangiectasia Type 2: A Randomized Clinical Trial. Ophthalmology, 2019. 126(4): p. 540–549.

32. Weinreb, R.N., et al., Primary open-angle glaucoma. Nat Rev Dis Primers, 2016. 2: p. 16067.

33. Ip, N.Y., et al., The alpha component of the CNTF receptor is required for signaling and defines potential CNTF targets in the adult and during development. Neuron, 1993. 10(1): p. 89–102.

34. Ip, N.Y., The neurotrophins and neuropoietic cytokines: two families of growth factors acting on neural and hematopoietic cells. Ann N Y Acad Sci, 1998. 840: p. 97–106.

35. Bonni, A., et al., Characterization of a pathway for ciliary neurotrophic factor signaling to the nucleus. Science, 1993. 262(5139): p. 1575–9.

36. Boulton, T.G., N. Stahl, and G.D. Yancopoulos, Ciliary neurotrophic factor/leukemia inhibitory factor/interleukin 6/oncostatin M family of cytokines induces tyrosine phosphorylation of a common set of proteins overlapping those induced by other cytokines and growth factors. J Biol Chem, 1994. 269(15): p. 11648–55.

37. Peterson, W.M., et al., Ciliary neurotrophic factor and stress stimuli activate the Jak-STAT pathway in retinal neurons and glia. J Neurosci, 2000. 20(11): p. 4081–90.

38. Rhee, K.D., et al., Cytokine-induced activation of signal transducer and activator of transcription in photoreceptor precursors regulates rod differentiation in the developing mouse retina. J Neurosci, 2004. 24(44): p. 9779–88.

39. Rhee, K.D., et al., Molecular and cellular alterations induced by sustained expression of ciliary neurotrophic factor in a mouse model of retinitis pigmentosa. Invest Ophthalmol Vis Sci, 2007. 48(3): p. 1389–400.

40. Rhee, K.D., et al., CNTF-mediated protection of photoreceptors requires initial activation of the cytokine receptor gp130 in Muller glial cells. Proc Natl Acad Sci U S A, 2013. 110(47): p. E4520–9.

41. Liang, F.Q., et al., Long-term protection of retinal structure but not function using RAAV.CNTF in animal models of retinitis pigmentosa. Mol Ther, 2001. 4(5): p. 461–72.

42. Bok, D., et al., Effects of adeno-associated virus-vectored ciliary neurotrophic factor on retinal structure and function in mice with a P216L rds/peripherin mutation. Exp Eye Res, 2002. 74(6): p. 719–35.

43. Wen, R., et al., Regulation of rod phototransduction machinery by ciliary neurotrophic factor. J Neurosci, 2006. 26(52): p. 13523–30.

44. Wang, Y., et al., Impacts of ciliary neurotrophic factor on the retinal transcriptome in a mouse model of photoreceptor degeneration. Sci Rep, 2020. 10(1): p. 6593.

45. Okawa, H., et al., ATP consumption by mammalian rod photoreceptors in darkness and in light. Curr Biol, 2008. 18(24): p. 1917–21.

46. Ingram, N.T., G.L. Fain, and A.P. Sampath, Elevated energy requirement of cone photoreceptors. Proc Natl Acad Sci U S A, 2020. 117(32): p. 19599–19603.

47. Hurley, J.B., Warburg's vision. Elife, 2017. 6.

48. Du, J., et al., Phototransduction Influences Metabolic Flux and Nucleotide Metabolism in Mouse Retina. J Biol Chem, 2016. 291(9): p. 4698–710.

49. Kanow, M.A., et al., Biochemical adaptations of the retina and retinal pigment epithelium support a metabolic ecosystem in the vertebrate eye. Elife, 2017. 6.

50. Hurley, J.B., Retina Metabolism and Metabolism in the Pigmented Epithelium: A Busy Intersection. Annu Rev Vis Sci, 2021.

51. Adijanto, J., et al., The retinal pigment epithelium utilizes fatty acids for ketogenesis. J Biol Chem, 2014. 289(30): p. 20570–82.

52. Reyes-Reveles, J., et al., Phagocytosis-dependent ketogenesis in retinal pigment epithelium. J Biol Chem, 2017. 292(19): p. 8038–8047.

53. Fu, Z., et al., Fatty acid oxidation and photoreceptor metabolic needs. J Lipid Res, 2021. 62: p. 100035.

54. Punzo, C., K. Kornacker, and C.L. Cepko, Stimulation of the insulin/mTOR pathway delays cone death in a mouse model of retinitis pigmentosa. Nat Neurosci, 2009. 12(1): p. 44–52.

55. Ait-Ali, N., et al., Rod-derived cone viability factor promotes cone survival by stimulating aerobic glycolysis. Cell, 2015. 161(4): p. 817–32.

56. Chertov, A.O., et al., Roles of glucose in photoreceptor survival. J Biol Chem, 2011. 286(40): p. 34700–11.

57. Hurley, J.B., K.J. Lindsay, and J. Du, Glucose, lactate, and shuttling of metabolites in vertebrate retinas. J Neurosci Res, 2015. 93(7): p. 1079–92.

58. Chinchore, Y., et al., Glycolytic reliance promotes anabolism in photoreceptors. Elife, 2017. 6.

59. Petit, L., et al., Aerobic Glycolysis Is Essential for Normal Rod Function and Controls Secondary Cone Death in Retinitis Pigmentosa. Cell Rep, 2018. 23(9): p. 2629–2642.

60. Zhang, R., et al., Selective knockdown of hexokinase 2 in rods leads to age-related photoreceptor degeneration and retinal metabolic remodeling. Cell Death Dis, 2020. 11(10): p. 885.

61. Rajala, A., et al., Pyruvate kinase M2 regulates photoreceptor structure, function, and viability. Cell Death Dis, 2018. 9(2): p. 240.

62. Rajala, A., et al., Pyruvate kinase M2 isoform deletion in cone photoreceptors results in age-related cone degeneration. Cell Death Dis, 2018. 9(7): p. 737.

63. Xu, L., et al., Stimulation of AMPK prevents degeneration of photoreceptors and the retinal pigment epithelium. Proc Natl Acad Sci U S A, 2018. 115(41): p. 10475–10480.

64. Kedzierski, W., et al., Deficiency of rds/peripherin causes photoreceptor death in mouse models of digenic and dominant retinitis pigmentosa. Proc Natl Acad Sci U S A, 2001. 98(14): p. 7718–23.

65. Li, S., et al., Rhodopsin-iCre transgenic mouse line for Cre-mediated rod-specific gene targeting. Genesis, 2005. 41(2): p. 73–80.

66. Pham, A.H., J.M. McCaffery, and D.C. Chan, Mouse lines with photo-activatable mitochondria to study mitochondrial dynamics. Genesis, 2012. 50(11): p. 833–43.

67. Liesa, M. and O.S. Shirihai, Mitochondrial dynamics in the regulation of nutrient utilization and energy expenditure. Cell Metab, 2013. 17(4): p. 491–506.

68. Vowinckel, J., et al., MitoLoc: A method for the simultaneous quantification of mitochondrial network morphology and membrane potential in single cells. Mitochondrion, 2015. 24: p. 77–86.

69. Guimaraes-Ferreira, L., Role of the phosphocreatine system on energetic homeostasis in skeletal and cardiac muscles. Einstein (Sao Paulo), 2014. 12(1): p. 126–31.

70. Chen, L., et al., In vivo imaging of phosphocreatine with artificial neural networks. Nat Commun, 2020. 11(1): p. 1072.

71. Malpartida, A.B., et al., Mitochondrial Dysfunction and Mitophagy in Parkinson's Disease: From Mechanism to Therapy. Trends Biochem Sci, 2021. 46(4): p. 329–343.

72. Ng, M.Y.W., T. Wai, and A. Simonsen, Quality control of the mitochondrion. Dev Cell, 2021. 56(7): p. 881–905.

73. Fu, Z., et al., Fatty acid oxidation and photoreceptor metabolic needs. J Lipid Res, 2020.

74. Noctor, G., et al., Glutathione: biosynthesis, metabolism and relationship to stress tolerance explored in transformed plants. Journal of Experimental Botany, 1998. 49(321): p. 623–647.

75. Warburg, O., F. Wind, and E. Negelein, The Metabolism of Tumors in the Body. J Gen Physiol, 1927. 8(6): p. 519–30.

76. Ly, C.H., G.S. Lynch, and J.G. Ryall, A Metabolic Roadmap for Somatic Stem Cell Fate. Cell Metab, 2020. 31(6): p. 1052–1067.

77. Tsogtbaatar, E., et al., Energy Metabolism Regulates Stem Cell Pluripotency. Front Cell Dev Biol, 2020. 8: p. 87.

78. Bromberg, J.F. and J.E. Darnell, Jr., Potential roles of Stat1 and Stat3 in cellular transformation. Cold Spring Harb Symp Quant Biol, 1999. 64: p. 425–8.

79. Bromberg, J. and J.E. Darnell, Jr., The role of STATs in transcriptional control and their impact on cellular function. Oncogene, 2000. 19(21): p. 2468–73.

80. Shen, Y., et al., Constitutively activated Stat3 protects fibroblasts from serum withdrawal and UV-induced apoptosis and antagonizes the proapoptotic effects of activated Stat1. Proc Natl Acad Sci U S A, 2001. 98(4): p. 1543–8.

81. Gough, D.J., et al., Mitochondrial STAT3 supports Ras-dependent oncogenic transformation. Science, 2009. 324(5935): p. 1713–6.

82. Wegrzyn, J., et al., Function of mitochondrial Stat3 in cellular respiration. Science, 2009. 323(5915): p. 793–7.

83. Szczepanek, K., et al., Mitochondrial-targeted Signal transducer and activator of transcription 3 (STAT3) protects against ischemia-induced changes in the electron transport chain and the generation of reactive oxygen species. J Biol Chem, 2011. 286(34): p. 29610–20.

84. Chowdhury, S.R., et al., Ciliary neurotrophic factor reverses aberrant mitochondrial bioenergetics through the JAK/STAT pathway in cultured sensory neurons derived from streptozotocin-induced diabetic rodents. Cell Mol Neurobiol, 2014. 34(5): p. 643–9.

85. Macias, E., et al., Stat3 binds to mtDNA and regulates mitochondrial gene expression in keratinocytes. J Invest Dermatol, 2014. 134(7): p. 1971–1980.

86. Jiang, K., et al., STAT3 promotes survival of mutant photoreceptors in inherited photoreceptor degeneration models. Proc Natl Acad Sci U S A, 2014. 111(52): p. E5716–23.

87. Conti, B., et al., Uncoupling protein 2 protects dopaminergic neurons from acute 1,2,3,6-methyl-phenyl-tetrahydropyridine toxicity. J Neurochem, 2005. 93(2): p. 493–501.

88. Andrews, Z.B., et al., Uncoupling protein-2 is critical for nigral dopamine cell survival in a mouse model of Parkinson's disease. J Neurosci, 2005. 25(1): p. 184–91.

89. Andrews, Z.B., S. Diano, and T.L. Horvath, Mitochondrial uncoupling proteins in the CNS: in support of function and survival. Nat Rev Neurosci, 2005. 6(11): p. 829–40.

90. Barnstable, C.J., et al., Mitochondrial Uncoupling Protein 2 (UCP2) Regulates Retinal Ganglion Cell Number and Survival. J Mol Neurosci, 2016. 58(4): p. 461–9.

91. Ou, J., et al., iPSCs from a Hibernator Provide a Platform for Studying Cold Adaptation and Its Potential Medical Applications. Cell, 2018. 173(4): p. 851–863 e16.

92. Flores, A., et al., Lactate dehydrogenase activity drives hair follicle stem cell activation. Nat Cell Biol, 2017. 19(9): p. 1017–1026.

93. Quansah, E., et al., Targeting energy metabolism via the mitochondrial pyruvate carrier as a novel approach to attenuate neurodegeneration. Mol Neurodegener, 2018. 13(1): p. 28.

94. Eade, K., et al., Serine biosynthesis defect due to haploinsufficiency of PHGDH causes retinal disease. Nat Metab, 2021. 3(3): p. 366–377.

95. Bonelli, R., et al., Identification of genetic factors influencing metabolic dysregulation and retinal support for MacTel, a retinal disorder. Commun Biol, 2021. 4(1): p. 274.

96. Bonelli, R., et al., Systemic lipid dysregulation is a risk factor for macular neurodegenerative disease. Sci Rep, 2020. 10(1): p. 12165.

97. Hashimoto, T., et al., Lentiviral gene replacement therapy of retinas in a mouse model for Usher syndrome type 1B. Gene Ther, 2007. 14(7): p. 584–94.

98. Wahlin, K.J., et al., A method for analysis of gene expression in isolated mouse photoreceptor and Muller cells. Mol Vis, 2004. 10: p. 366–75.

99. Jin, M., et al., The role of interphotoreceptor retinoid-binding protein on the translocation of visual retinoids and function of cone photoreceptors. J Neurosci, 2009. 29(5): p. 1486–95.

100. Schneider, C.A., W.S. Rasband, and K.W. Eliceiri, NIH Image to ImageJ: 25 years of image analysis. Nat Methods, 2012. 9(7): p. 671–5.

101. Linkert, M., et al., Metadata matters: access to image data in the real world. J Cell Biol, 2010. 189(5): p. 777–82.

102. Otsu, N., A threshhold selection method from gray-level histograms. IEEE Transactions on Systems, Man, and Cybernetics, 1979. 9(1): p. 62–66.

103. Sebastien Le, J.J., Husson, F., An R package for multivariable analysis. Journal of Statistical Software, 2008. 25(1): p. 1–18.

104. Kooragayala, K., et al., Quantification of Oxygen Consumption in Retina Ex Vivo Demonstrates Limited Reserve Capacity of Photoreceptor Mitochondria. Invest Ophthalmol Vis Sci, 2015. 56(13): p. 8428–36.

105. Pertea, M., et al., Transcript-level expression analysis of RNA-seq experiments with HISAT, StringTie and Ballgown. Nat Protoc, 2016. 11(9): p. 1650–67.

